# Novel entropy-based metrics for predicting choice behavior based on local response to reward

**DOI:** 10.1101/2021.05.20.445009

**Authors:** Ethan Trepka, Mehran Spitmaan, Bilal A. Bari, Vincent D. Costa, Jeremiah Y. Cohen, Alireza Soltani

**Affiliations:** Department of Psychological and Brain Sciences, Dartmouth College, NH, USA; The Solomon H. Snyder Department of Neuroscience, Brain Science Institute, Kavli Neuroscience Discovery Institute, The Johns Hopkins University School of Medicine, MD, USA; Department of Behavioral Neuroscience, Oregon Health and Science University, OR, USA

**Keywords:** Reinforcement learning, information theory, matching law, win-stay, lose-switch

## Abstract

For decades, behavioral scientists have used the matching law to quantify how animals distribute their choices between multiple options in response to reinforcement they receive. More recently, many reinforcement learning (RL) models have been developed to explain choice by integrating reward feedback over time. Despite reasonable success of RL models in capturing choice on a trial-by-trial basis, these models cannot capture variability in matching. To address this, we developed novel metrics based on information theory and applied them to choice data from dynamic learning tasks in mice and monkeys. We found that a single entropy-based metric can explain 50% and 41% of variance in matching in mice and monkeys, respectively. We then used limitations of existing RL models in capturing entropy-based metrics to construct a more accurate model of choice. Together, our novel entropy-based metrics provide a powerful, model-free tool to predict adaptive choice behavior and reveal underlying neural mechanisms.

## Introduction

How do we distribute our time and choices between the many options or actions available to us? Around 60 years ago, Richard Herrnstein strived to answer this question based on one of the main ideas of behaviorism; that is, the history of reinforcement is the most important determinant of behavior. He proposed a simple rule called the matching law stating that the proportion of time or responses that an animal allocates to an option or action matches the proportion of reinforcement they receive from those options or actions (Herrnstein, 1961). The matching law has been shown to explain global choice behavior across many species (Williams, 1988) including pigeons (de Villiers and Herrnstein, 1976; William, 1979; Mazur, 1981; Villarreal et al., 2019), mice (Gallistel et al., 2007; Fonseca et al., 2015; Bari et al., 2019), rats (Gallistel, 1994; Belke and Belliveau, 2001; Lee et al., 2017), monkeys (Anderson et al., 2002; Sugrue et al., 2004; Lau and Glimcher, 2005; Kubanek and Snyder, 2015; Tsutsui et al., 2016; Soltani et al., 2021), and humans (Schroeder and Holland, 1969; Pierce and Epling, 1983; Beardsley and McDowell, 1992; Savastano and Fantino, 1994; Vullings and Madelain, 2018; Cero and Falligant, 2020), in a wide range of tasks, including concurrent variable interval, concurrent variable ratio, probabilistic reversal learning, and so forth. A common finding in most studies, however, has been that animals undermatch, corresponding to selection of the better option/action less than it is prescribed by the matching law. Such deviation from matching often corresponds to suboptimal behavior in terms of total harvested rewards, pointing to adaptive mechanisms beyond reward maximization.

The matching law is a global rule but ultimately should emerge from an interaction between choice and learning strategies, resulting in local (in time) adjustments of choice behavior to reinforcement obtained in each trial. Accordingly, many studies have tried to explain matching based on different mechanisms. Reinforcement learning models are particularly useful because they can simulate changes in behavior due to reward feedback. Consequently, recent studies on matching have focused on developing RL models that can generate global matching behavior based on local learning rules (Sugrue et al., 2004; Lau and Glimcher, 2005; Loewenstein and Seung, 2006; Soltani and Wang, 2006; Ito and Doya, 2009; Otto et al., 2011; Iigaya and Fusi, 2013; Saito et al., 2014; Iigaya et al., 2019; Grossman et al., 2020). These RL models are often augmented with some components in addition to stimulus- or action-value functions to improve fit of choice behavior on a trial-by-trial basis. For example, the models could include learning the reward-independent rate of choosing each option (Lau and Glimcher, 2005), adopting win-stay lose-switch policies (Ito and Doya, 2009; Otto et al., 2011), or learning on multiple timescales (Iigaya et al., 2019). Although these additional components improved the overall fit of choice behavior, the outcome models were not examined for their ability to capture deviation from matching. This could result in misinterpretation or missing important neural mechanisms underlying matching behavior in particular and adaptive behavior more generally (Palminteri et al., 2017; Wilson and Collins, 2019). Therefore, after decades of research on matching behavior, it is still not fully clear how such a fundamental law of behavior emerges as a result of local response to reward feedback.

To address this, we propose a set of novel metrics based on information theory that can summarize trial-by-trial response to reward feedback and predict global matching behavior. To test the utility of our novel metrics, we applied them to large sets of behavioral data in mice and monkeys during two very different dynamic learning tasks. We found that in both mice and monkeys, our entropy-based metrics can predict matching behavior better than existing measures. Moreover, we found a close link between undermatching and the consistency of choice strategy (stay or switch) in response to receiving no reward after selection of the worse option in both species. Finally, we used shortcomings of existing RL models in capturing the pattern of entropy-based metrics in our data to construct a new model that integrates reward- and option-dependent strategies with a reinforcement-learning component. We show that the new model can capture both trial-by-trial choice data and global choice behavior better than the existing models.

## Results

### Mice and monkeys dynamically adjusted their behavior to changes in reward probabilities

To study learning and decision making in dynamic reward environments, we examined choice behavior of mice and monkeys during two different probabilistic reversal learning tasks. Mice selected between two actions (licking left and right) that provided reward with different probabilities, and these probabilities changed between blocks of trials without any signal to the animals (**Fig. 1a**; see **Methods** for more details; Bari et al., 2019). Block lengths were drawn from a uniform distribution that spanned a range of 40 to 80 trials. Here, we focused on the majority (469 out of 528) of sessions in which two sets of reward probabilities (equal to 0.4 and 0.1, and 0.4 and 0.05) were used. We refer to these reward schedules as 40/10 and 40/5 reward schedules (1786 and 1533 blocks with 40/5 and 40/10 reward schedules, respectively). Rewards were “baited” such that if reward was assigned on a given side and that side was not selected, reward would remain on that side until the next time that side was selected. Due to baiting, the probability of obtaining reward on the unchosen side increased over time as during foraging in a natural environment. As a result, optimal performance in the task generally required a strategy that involved selecting options in proportion to their reinforcement rates (perfect “matching” behavior). In total, 16 mice performed 469 sessions of the two-probability version of the task for a total of 3319 blocks and 189,199 trials.

**Figure 1.**
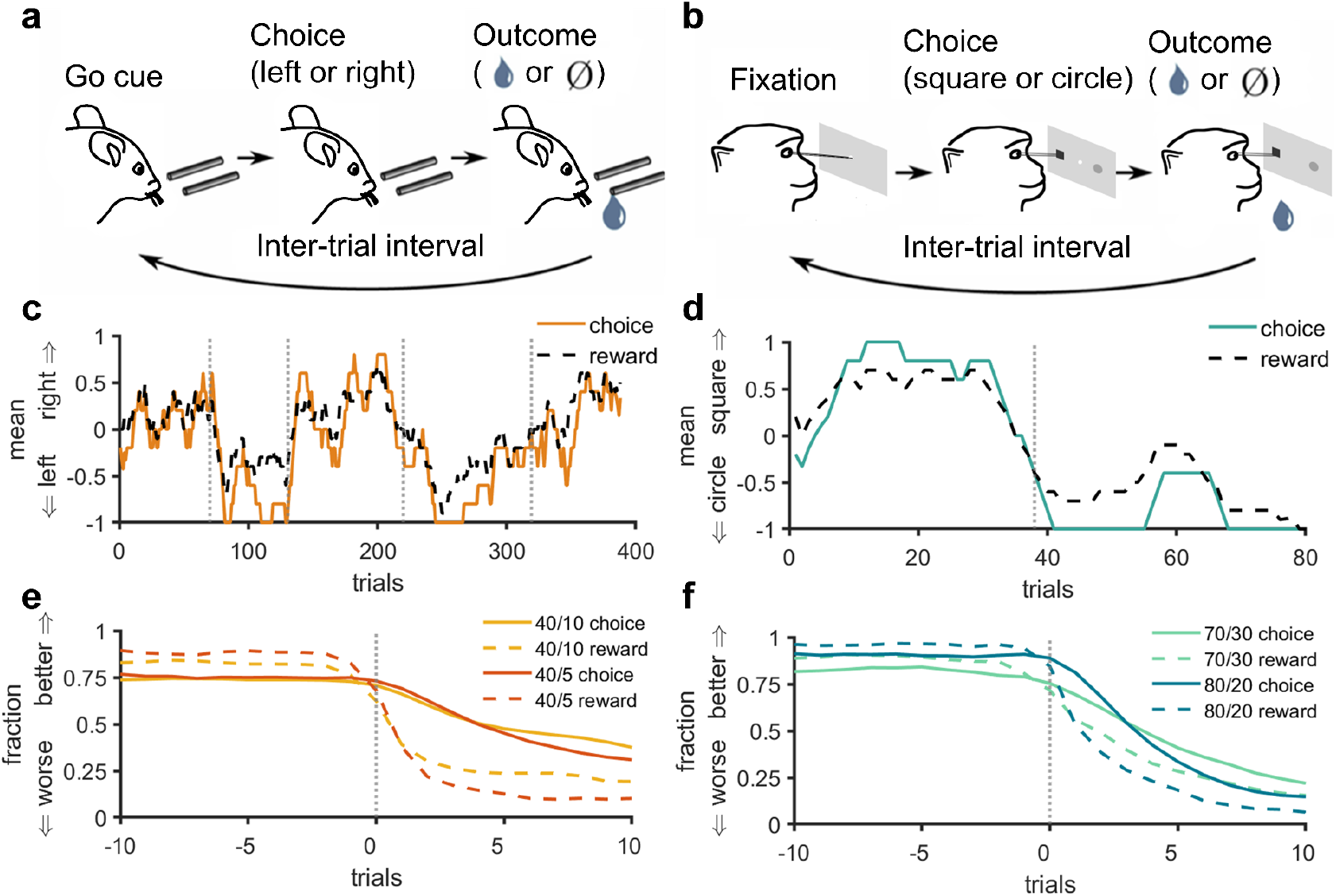
Schematic of the experimental paradigms in mice and monkeys and basic behavioral results. **(a–b)** Timeline of a single trial during experiments in mice (a) and monkeys (b). To initiate a trial, mice received an olfactory “go” cue (or “no go” cue in 5% of trials) (a), and monkeys fixated on a central point (b). Next, animals chose (via licks for mice and saccades for monkeys) between two options (left or right tubes for mice and circle or square for monkeys) and then received a reward (drop of water and juice for mice and monkeys, respectively) probabilistically based on their choice. **(c–d)** Average choice and reward using a sliding window with a length of 10 for two representative sessions in mice (c) and monkeys (d). Grey dashed lines indicate trials where reward probabilities reversed. **(e–f)** Average choice and reward fractions around block switches using a non-causal smoothing kernel with a length of 3 separately for all blocks with a given reward schedule in mice (e) and monkeys (f). The better (or worse) side is the better (or worse) side prior to the block switch. Trial zero is the first trial with the reversed reward probabilities. Average choice fractions for the better side/stimulus are lower than average reward fractions for that side/stimulus throughout the block for both mice and monkeys, corresponding to undermatching behavior.

In a different experiment, monkeys selected between two new stimuli (a circle or square with variable colors) via saccades and received a juice reward probabilistically (**Fig. 1b**; see **Methods** for more details; Costa et al., 2016). In superblocks of 80 trials, the reward probabilities assigned to each stimulus reversed randomly between trials 30 and 50, such that the more rewarding stimulus became the less rewarding stimulus. We refer to trials before and after a reversal as a block. Monkeys completed multiple superblocks per each session of the experiment wherein the reward probabilities assigned to the better and worse options were equal to 0.8 and 0.2, 0.7 and 0.3, or 0.6 and 0.4, which we refer to as 80/20, 70/30, and 60/40 reward schedules. In contrast to the task used in mice, rewards were not baited. Here, we only analyze data from the 80/20 and 70/30 reward schedules as they provide two levels of reward uncertainty similar to the experiment in mice. In total, 4 monkeys performed 2212 blocks of the task with the 80/20 and 70/30 reward schedules for a total of 88,480 trials.

We found that in response to block switches, both mice and monkeys rapidly adjusted their choice behavior to select the better side or stimulus more often (**Fig. 1c–d**). However, the fraction of times they chose the better side or stimulus fell below predictions made by the matching law, even at the end of the blocks (**Fig. 1e–f**). More specifically, the relative selection of the better side/stimulus (choice fraction) was often lower than the ratio of reward harvested on the better side/stimulus to the overall reward harvested (reward fraction), corresponding to undermatching behavior. Therefore, we next explored how undermatching depends on choice- and reward-dependent strategies.

### Mice and monkeys exhibited highly variable undermatching behavior

To better examine matching behavior, we used the difference between the relative choice and reward fractions for each block of trials to define “deviation from matching” (see **Eqs. 1–2** in **Methods**; **Fig. 2a,d**). Based on our definition, negative and positive values for deviation from matching correspond to undermatching and overmatching, respectively. Undermatching occurs when the choice fraction is smaller than the reward fraction for reward fractions larger than 0.5, or the choice fraction is larger than the reward fraction when the latter is smaller than 0.5. Overmatching occurs when the choice fraction is larger than the reward fraction for reward fractions larger than 0.5, or the choice fraction is smaller than the reward fraction when the latter is smaller than 0.5. Undermatching could happen because the animal does not detect the more rewarding option/action, poor credit assignment, or due to too much stochasticity in choice. In contrast, overmatching is characterized by selecting the better option more frequently than is prescribed based on perfect matching. In the task used in monkeys, overmatching was not possible by design (except due to random fluctuations in reward assignment) and optimal performance could be achieved by selecting the better stimulus all the time, corresponding to matching. In contrast, while being sub-optimal, overmatching is possible in the reversal learning tasks with baited rewards (e.g., task used in mice).

**Figure 2.**
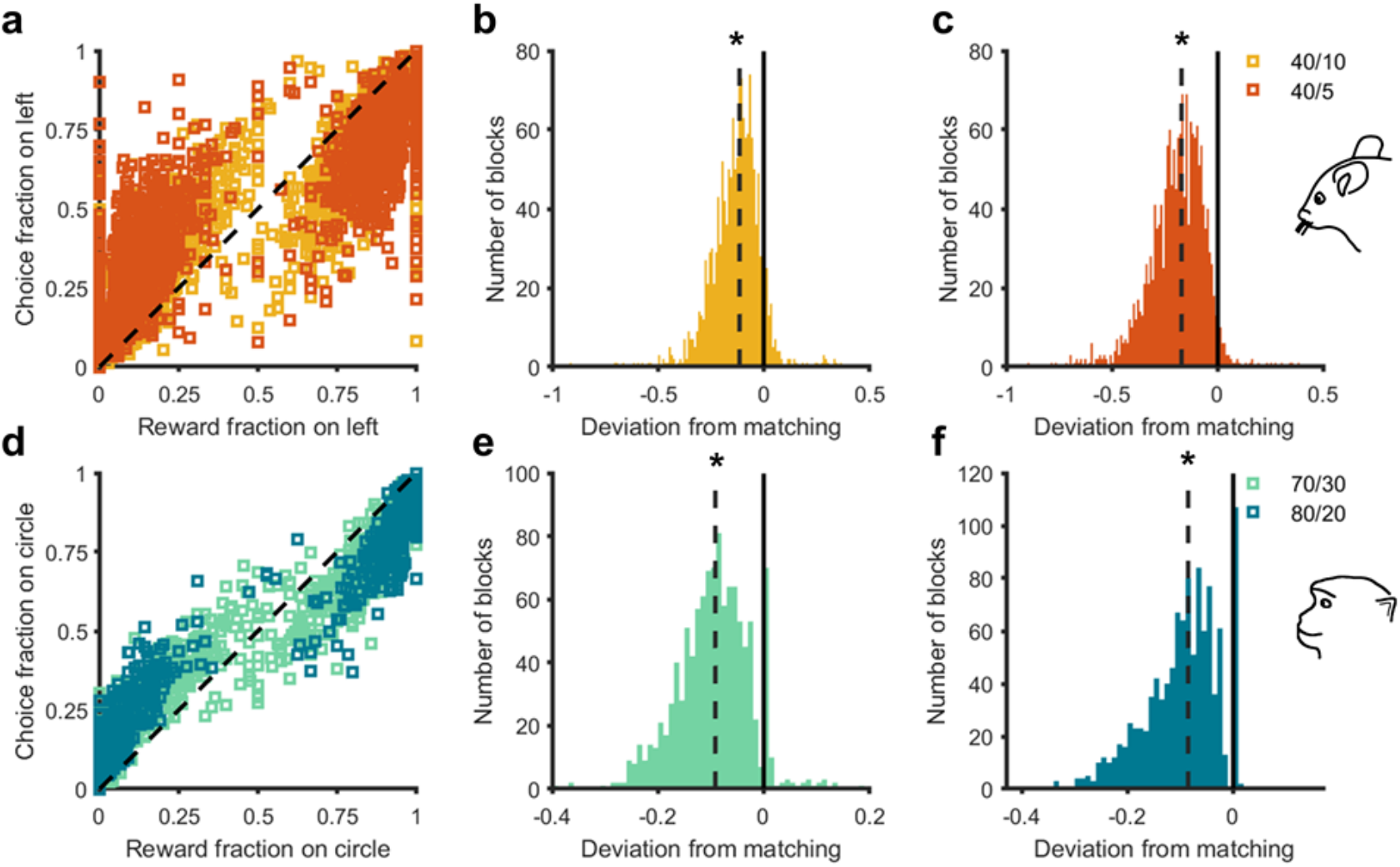
Mice and monkeys exhibited highly variable but significant undermatching. **(a)** Plot shows reward fraction versus choice fraction across all blocks separately for each reward schedule in mice. The black dashed line represents the identity line. For reward fractions greater than 0.5, nearly all points remain below the identity line, indicating choice fraction smaller than reward fraction for the better option (undermatching). Similarly, points above the identity line for reward fractions less than 0.5 indicate undermatching. (**b–c**) Distributions show deviation from matching values for the 40/10 (b) and 40/5 reward schedules (c) in mice. The solid black line indicates 0, corresponding to perfect matching. The dashed black lines are the median deviation from matching. Asterisks indicate significance (*p* < 0.01) based on a two-sided Wilcoxon signed-rank test. (**d–f**) Similar to a-c, but for monkeys with 70/30 and 80/20 reward schedules. Because of random fluctuations in local reward probabilities, overmatching occurred in a minority of cases.

Consistent with previous studies on matching behavior, we found significant undermatching in mice in both the 40/10 and 40/5 reward schedules (Wilcoxon signed-rank test; 40/10: *Z* = −31.2,*p* = 1.53 × 10^-213^; 40/5: *Z* = −35.0,*p* = 8.75 × 10^-269^; **Fig. 2b-c**). Similarly, we found significant undermatching in monkeys in both the 70/30 and 80/20 reward schedules (Wilcoxon signed-rank test; 70/30: *Z* = −27.02, *p* = 9.74 × 10^-161^; 80/20: *Z* = −27.06, *p* = 3.06 × 10^-161^; **Fig. 2e-f**). In addition, average undermatching for mice was significantly larger in the 40/5 reward schedule than the 40/10 reward schedule, whereas average undermatching for monkeys was not significantly different in the 70/30 and 80/20 schedules (two-sided independent t-test; Mice: *p* = 7.93 × 10^-36^, *d* = 0.44; Monkeys: *p* = 0.19, d = 0.06; **Fig. S1f**). More importantly, undermatching was highly variable in both reward schedules for both mice and monkeys (**Fig. 2b-c, 2e-f**). To understand the nature of this variability, we examined whether existing behavioral metrics and RL models can predict the observed deviation from matching.

### Existing metrics only partially explain variability in undermatching behavior

To examine the relationship between existing behavioral metrics and undermatching, we first performed stepwise multiple regressions to predict deviation from matching for both mice and monkeys based on commonly used metrics including *p*(*win*), *p*(*stay*), *p*(*stay|win*) and *p*(*switch|lose*) (threshold for adding a predictor: p<0.0001; see **Methods** for more details). These stepwise regressions resulted in the following equations for predicting deviation from matching in mice and monkeys:

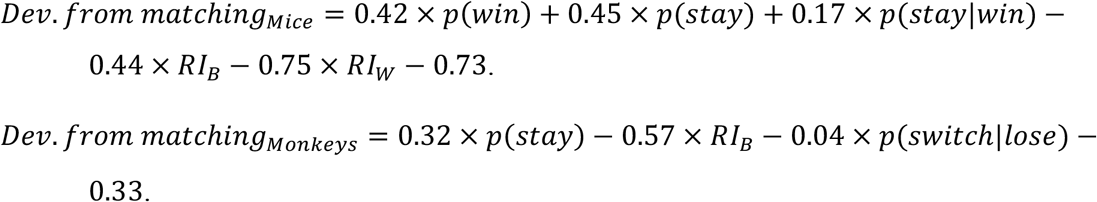

These regression models explained 31% and 34% of the variance in deviation from matching for mice and monkeys, respectively, which are significant but unsurprising amounts of overall variance (Mice: *Adjusted R*^2^ = 0.31; Monkeys: *Adjusted R*^2^ = 0.34). In the final regression model, *p*(*win*) and *p* (*stay|win*) all had positive regression coefficients for both mice and monkeys, indicating that increased *p*(*win*) and *p*(*stay|win*) are associated with less undermatching behavior (note that deviation from matching is mostly negative).

We next included the Repetition Index (RI) on the better (RI_B_) and worse (RI_W_) options (side or stimulus), which measure the tendency to stay beyond chance on the better and worse options, to predict undermatching (Soltani et al., 2013). To that end, we conducted additional stepwise multiple regressions that predicted deviation from matching using *RI_B_, RI_W_, p*(*win*), *p*(*stay*), *p*(*stay|win*), and *p*(*switch|lose*) as predictors. The final regression equations for predicting deviation from matching for mice and monkeys were:

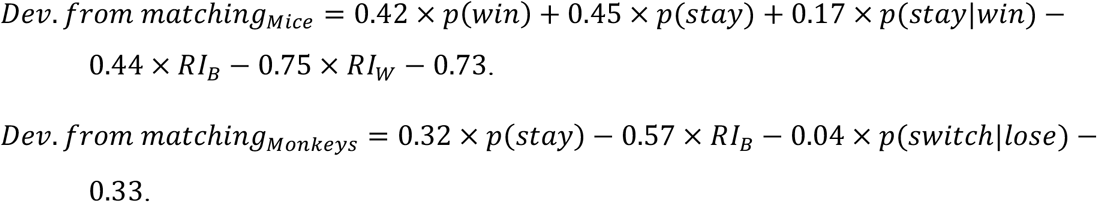

These models explained 48% and 49% of the variance in deviation from matching for mice and monkeys, respectively (Mice: *Adjusted R*^2^ = 0.48; Monkeys: *Adjusted R*^2^ = 0.49). Thus, including RI_B_ and RI_W_ enabled us to account for 17% and 15% more variance, suggesting that staying beyond chance on both the better and worse sides is a significant contributor to undermatching behavior. RI_B_ and RI_W_ both had negative coefficients in the regression equations they were present in, indicating that repeating better or worse options beyond chance increases undermatching. This was expected for RI_W_ because staying beyond chance on the worse side results in more frequent selection of the worse option and thus more undermatching. In contrast, larger RI_B_ could increase undermatching because more staying on the better side could stop the animals from switching to the new better side after block switches.

Together, these results illustrate that undermatching is correlated with tendency to stay beyond chance (measured by RI) and response to reward feedback in terms of stay or switch (measured by win-stay and lose-switch). However, win-stay and lose-switch are not strong predictors of undermatching because their relative importance depends on the overall probability of winning. For example, if overall win rate is high, lose-switch is less useful for predicting behavior because response to loss represents strategy in a small subset of trials. Although win-stay, lose-switch, *p*(*win*), and p(stay) contain all the information necessary to compute the dependence of strategy on reward, this requires interpretation of all four metrics in conjunction and may depend on nonlinear relationships that are challenging to intuit or capture with regression. To overcome these issues, we propose new metrics to quantify changes in strategy due to reward outcome using information theory.

### Novel behavioral metrics based on information theory

To better capture the dependence of staying (or similarly switching) strategy on reward outcome, we developed a series of model-free behavioral metrics based on Shannon’s information entropy (Shannon, 1948). The Shannon information entropy of a random variable X conditioned on Y, denoted H(X|Y), captures the surprise or uncertainty of X given knowledge of the values of Y. Lower information entropies correspond with decreased uncertainty in the variable under consideration and thus consistency in utilized strategy (see below).

First, we define the entropy of reward-dependent strategy (ERDS) that measures the dependence of staying and switching strategy on reward. Formally, ERDS is the information entropy of strategy conditioned on reward, *H*(*str|rew*) (see **Methods** for more details). ERDS quantifies the amount of information needed to explain an animal’s strategy given knowledge of whether the animal won or lost in the previous trial. Lower ERDS values indicate more consistent response to reward feedback.

In its simplest formulation for measuring the effect of reward in the previous trial on staying or switching, ERDS is a function of win-stay, lose-switch, and *p*(*win*) (see Eq. 9 in **Methods**). As win-stay and lose-switch move further from 0.5, ERDS decreases, reflecting increased consistency of reward-dependent strategy (**Fig. 3a)**. Moreover, *p*(*win*) modulates the effects of win-stay and lose-switch on ERDS (**Fig. 3b–c)**. As *p*(*win*) decreases, the influence of win-stay on ERDS decreases, reflecting that win-stay is less relevant to overall response to reward feedback when winning rarely occurs. Similarly, as *p*(*lose*) (=1-*p*(*win*)) decreases, the influence of lose-switch on ERDS decreases, reflecting that lose-switch is less relevant to response to reward feedback. Because of these properties, ERDS corrects for the limitations of win-stay and lose-switch. Also, as staying (or switching) strategy becomes more independent of reward outcome, ERDS increases because reward outcome provides no information about strategy.

**Figure 3.**
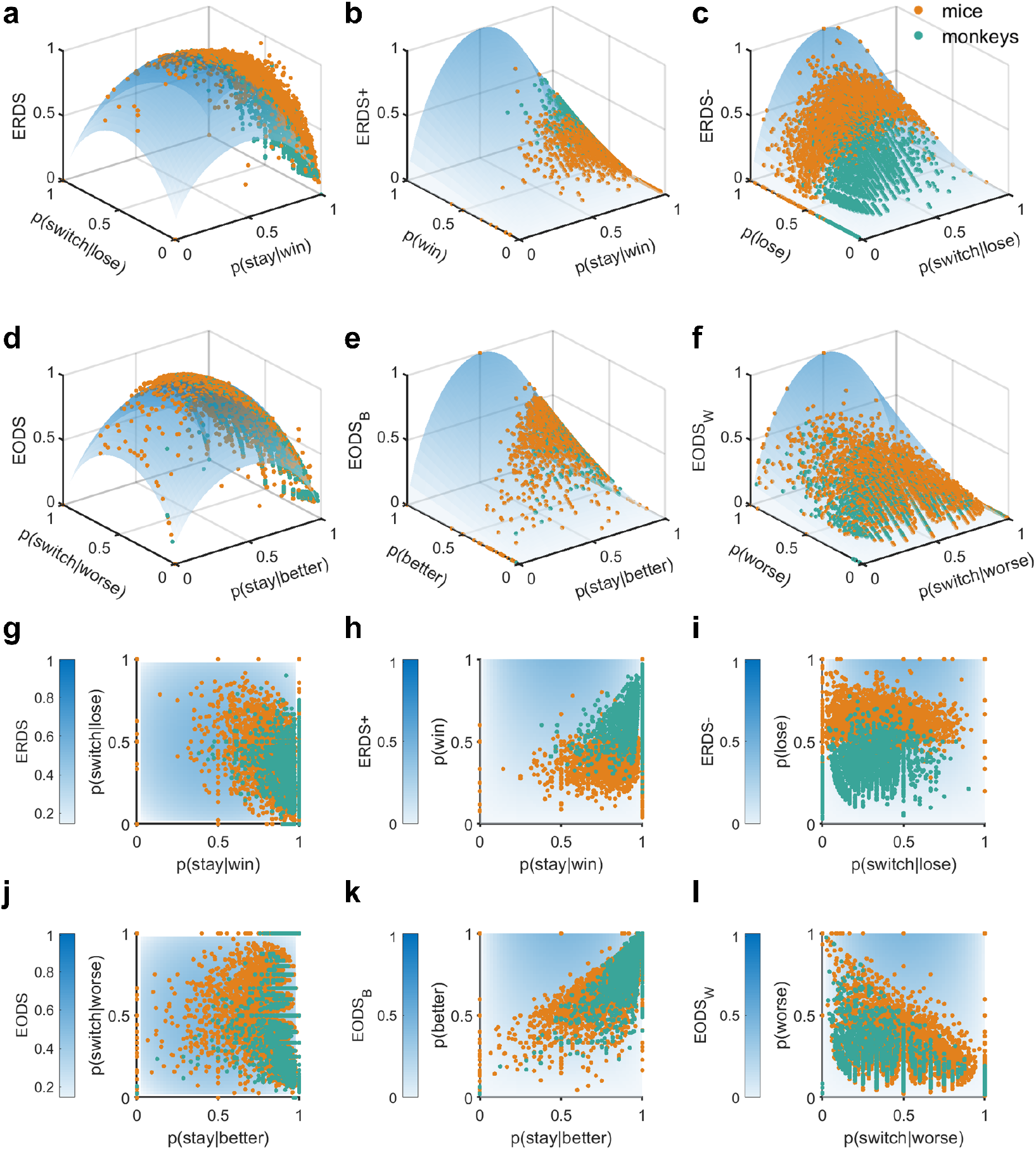
Relationship between new entropy-based metrics and win-stay, lose-switch strategies. (**a–c**) Plotted are ERDS and ERDS decompositions as a function of *p*(*win*), p(lose), win-stay, and lose-switch. Darker colors correspond to larger values of metrics. For the plot in panel A, *p*(*win*) is set to 0.5. Observed entropy-based metrics and constituent probabilities for each block for mice (orange dots) and monkeys (green dots) are superimposed on surfaces. (**d–f**) EODS and EODS decompositions as a function of the probabilities of choosing the better and worse option, p(better) and p(worse), probability of stay on the better option, and probability of switch from the worse option. For the plot in panel (d), p(better) is set to 0.5. For all plots, the units of entropy-based metrics are bits. (**g–i**) Same as in panels (a–c) but using heatmap. (**j–l**) Same as in panels (d-f) but using heatmap.

ERDS can be decomposed into ERDS+ and ERDS- to measure the specific effects of winning and losing in the preceding trial, respectively (**Fig. 3b-c;** see **Methods**). More specifically, ERDS+ is the entropy of win-dependent strategy, and ERDS- is the entropy of loss-dependent strategy. Therefore, comparing ERDS+ and ERDS- provides information about the relative contributions of win-dependent strategy and loss-dependent strategy to overall reward-dependent strategy.

In addition to conditioning stay strategy on reward in the preceding trial, we can also condition stay strategy on selection of the better or worse option. The resulting entropy of option-dependent strategy (EODS), *H*(*str|opt*), captures the dependence of staying and switching strategy on the selection of the better or worse option (or action) in the preceding trial (**Fig. 3d)**. EODS depends on p(choose better),*p*(*stay|choose better*), and *p*(*switch|choose worse*) and moreover, can be decomposed into EODS_B_ and EODS_W_ based on selection of the better or worse option (or action), respectively (**Fig. 3e-f)**.

Finally, to capture the dependence of strategy on reward and option in a single metric, we computed the entropy of reward- and option-dependent strategy (ERODS), *H*(*str|rew, opt*). ERODS depends on the probabilities of staying and switching conditioned on all combinations of reward and option (see **Methods** for more details). ERODS has similar properties to ERDS and EODS and can be interpreted in a similar fashion. Low ERODS values indicate that staying and switching strategy consistently depends on combinations of reward and the option selected in the previous trial; for example, winning and choosing the better side in the previous trial. ERODS can be decomposed either by better and worse option (ERODS_B_ and ERODS_W_), by win or loss (ERODS+ and ERODS-), or by both (ERODS_B_+, ERODS_W_+, ERODS_B_-, and ERODS_W−_).

To summarize, we propose three novel metrics, ERDS, EODS, and ERODS that capture the dependence of stay/switch strategy on reward outcome and/or selected option in the preceding trial. Each metric can be decomposed into components that provide important information about the dependence of strategy on winning or losing and/or choosing the better or worse option in the preceding trial (see **Table S1** for summary). We next show how these entropy-based metrics can predict deviation from matching behavior and further be used to construct more successful RL models.

### Deviation from matching was highly correlated with entropy-based metrics

To test the relationship between the observed undermatching and our novel entropy-based metrics, we next computed correlations between all behavioral metrics and deviation from matching (**Fig. 4)**. We found that nearly all entropy-based metrics were significantly correlated with deviation from matching. Out of all behavioral metrics tested, deviation from matching showed the strongest correlation with ERODS_W−_ based on both parametric (*Pearson; mice: r* = −0.71, *p* < 10^-300^; *monkeys: r* = −0.64, *p* = 10^-231^) and non-parametric (*Spearman; mice: r* = −0.78, *p* < 10^-300^; *monkeys: r* = −0.75, *p* < 10^-300^) tests in both mice and monkeys. The size of the correlation between ERODS_W−_ and deviation from matching is remarkable because it indicates that a single metric can capture more than 50% and 41% of the variance in deviation from matching in mice and monkeys, respectively. This finding suggests that undermatching occurs when animals lose when selecting the worse option and respond inconsistently to those losses.

**Figure 4.**
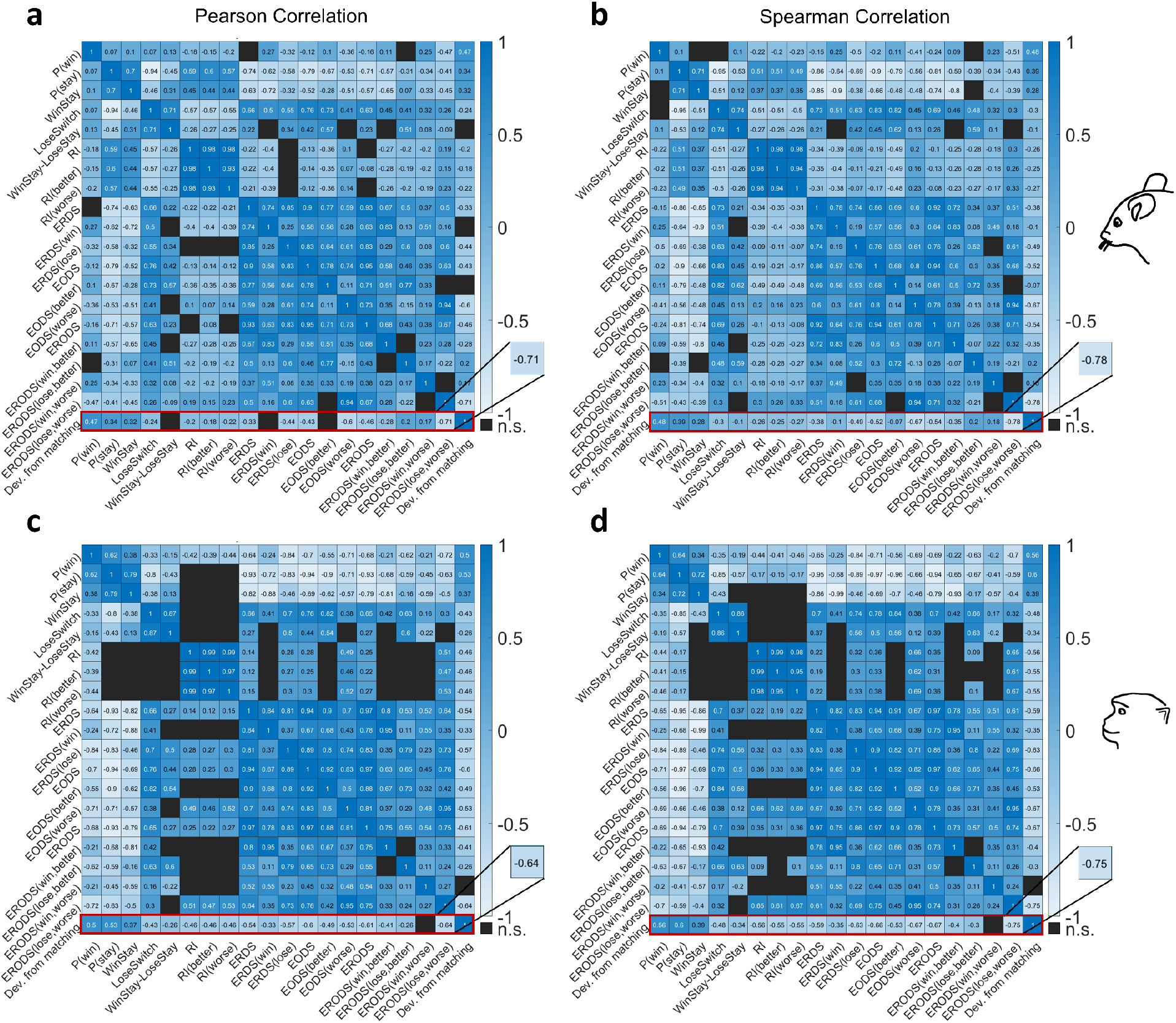
Correlation between undermatching and proposed entropy-based metrics and underlying probabilities. **(a-b)** Correlation matrix for 19 behavioral metrics and undermatching in mice using Pearson (a) and Spearman (b) tests. Correlation coefficients are computed across all blocks, and matrix elements with non-significant values (*p* > .0001) are not shown (cells in black). The red rectangles highlight correlation coefficients between behavioral metrics and undermatching. **(c-d)** Similar to (a-b) but for monkeys. Overall, the new entropy-based metrics show stronger correlation with undermatching than previous metrics, and undermatching was most strongly correlated with ERODS_W−_, EODS_W_, and ERDS-.

In addition to ERODS_W−_, EODS_W_ was also highly correlated with deviation from matching (*Pearson: mice: r* = −0.60, *p* < 10^-300^; *monkeys: r* = −0.53, *p* = 9.35 × 10^-150^; Spearman: mice: r = −0.67, *p* < 10^-300^, *monkeys: r* = −0.67 *p* = 1.18 × 10^-265^) as was ERDS- (*Pearson: mice: r* = −0.43, *p* = 6.72 × 10^-156^; *monkeys: r* = −0.57, *p* = 1.94 × 10^-191^; *Spearman: mice: r* = −0.47, *p* = 7.31 × 10^-198^; *monkeys: r* = −0.63, *p* = 1.06 × 10^-240^). Overall, these results show that global deviation from matching was most strongly correlated with the consistency of response after selection of the worse option (or action) and when no reward was obtained.

As expected, there were significant correlations between the proposed entropy-based metrics (see lower right of matrices in **Fig. 4**). For instance, ERDS and EODS were highly correlated (*Pearson: mice: r* = 0.90, *p* < 10^-300^; *monkeys: r* = 0.94, *p* < 10^-300^). EODS and ERODS were also highly correlated as expected (*Pearson: mice: r* = 0.95, *p* < 10^-300^; *monkeys: r* = 0.97, *p* < 10^-300^). Many entropy-based metric decompositions had similarly large correlations with other entropy-based metric decompositions. Finally, we found consistent results for the relationships between undermatching and our metrics for both reward schedules, even though undermatching and our metrics were sensitive to reward probabilities on the two options (see **Fig. S1**, **Fig. S2**, **Fig. S3**, and **Fig. S4**).

### Entropy-based metrics can accurately predict deviation from matching

To verify the utility of entropy-based metrics in predicting deviation from matching, we performed additional stepwise regressions using our entropy-based metrics. In these models, we included *ERDS_+_, ERDS_−_, EODS_B_, EODS_W_, ERODS_B+_, ERODS_B−_, ERODS_W+_, ERODS_W−_, RI_B_, RI_W_*, *p*(*win*), *p*(*stay*), *win-stay*, and *lose-switch* as predictors.

The final regression equations for predicting undermatching using all metrics in mice and monkeys were as follows:

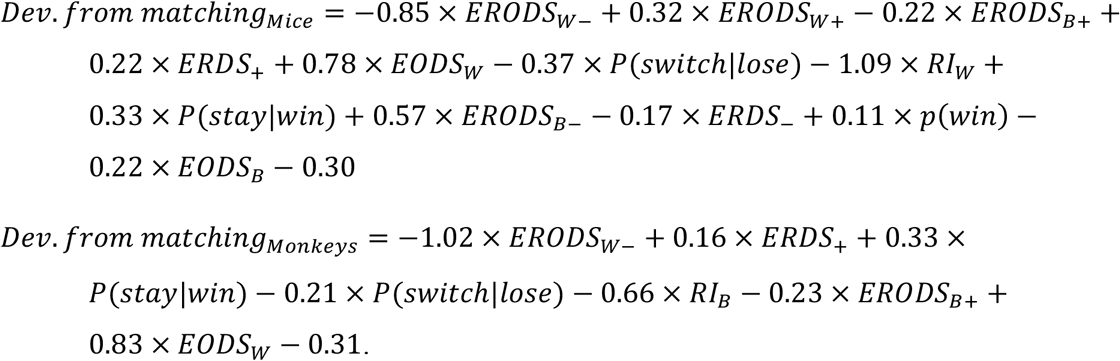

These models explained 74% and 57% of total variance in deviation from matching for mice and monkeys, respectively (Mice: *Adjusted R*^2^ = 0.74; Monkeys: *Adjusted R*^2^ = 0.57). For mice, the regression model explained 26% more variance than the model with repetition indices and 43% more variance than the model with basic behavioral metrics. For monkeys, the regression model explained 8% more variance than the model with repetition indices and 23% more variance than the model with basic behavioral metrics. These are significant improvements over previous models, suggesting that most variance in undermatching behavior can be explained by trial-by-trial response to reward feedback. Moreover, for mice, the first three predictors added to the regression models were ERODS_W−_ (Δ*R*^2^ = 0.59), ERODS_W_+ (Δ*R*^2^ = 0.04), and ERODS_B_+ (Δ*R*^2^ = 0.02). For monkeys, the first three predictors added were ERODS_W−_ (Δ*R*^2^ = 0.31), EODS_W_ (Δ*R*^2^ = 0.09), and ERODS_B_+ (Δ*R*^2^ = 0.06). Notably, the first three predictors added to the regression models for both mice and monkeys were entropy-based metrics, indicating that entropy-based metrics were the best predictors of deviation from matching when considering all metrics together.

The coefficients of predictors in this model cannot be interpreted in isolation in this model due to multicollinearity among entropy-based metrics. However, seven out of eight and four out of eight of the entropy-based metrics included as predictors were added to the final regression equations for mice and monkeys, respectively. This suggests that despite their overlap, each entropy-based metric captures a unique aspect of the variance in deviation from matching behavior.

Given the complexity of the final equation to predict deviation from matching, we also constructed simpler linear regression models predicting deviation from matching using the first three entropy-based metrics added to the stepwise regressions (*ERODS_W−_, ERODS_W+_*, and *ERODS_B+_* for mice, and *ERODS_W−_, ERODS_B+_*, and for monkeys). For mice, the regression equation for this simple model was: *Dev. from matching* = −0.62 × *ERODS_W−_* + 0.73 × *ERODS_W+_* — 0.16 × *ERODS_B+_* — 0.02. For monkeys, the regression equation for this simple model was: *Dev. from matching* = −1.09 × *ERODS_W−_* – 0.14 × *ERODS_B+_* + 0.63 × *EODS_W_* – 0.06. Despite these models’ simplicity, they still explained 64% of total variance in deviation from matching for mice and 45% for monkeys (Monkeys: *Adjusted R*^2^ = 0.45; Mice: *Adjusted R*^2^ = 0.64).

### Entropy-based metrics capture the relationship between undermatching and reward environment better than existing metrics

Our previous observation that entropy-based metrics can explain most variance in undermatching behavior suggests that entropy-based metrics may also capture average differences in undermatching between reward schedules in mice and the lack of differences in undermatching between reward schedules in monkeys.

To test whether this was the case, we used 10-fold cross validated linear regression to predict deviation from matching using a model without entropy-based or repetition metrics, a model without repetition metrics, and a model with all metrics. Predictors chosen for inclusion in these models were the predictors that remained in final stepwise regression equations described above. For mice, in all three models, predicted deviation from matching was significantly lower in the 40/5 than the 40/10 reward environment (**Fig. S5**; two-sided t-test; Model without entropy/repetition metrics: *p* = 6.30 × 10^-3^, Model without entropy-based metrics: *p* = 3.22 × 10^-3^, Full model with entropy and repetition metrics: *p* = 4.76 × 10^-25^). Importantly, the difference between deviation from matching in the two reward schedules was greatest for the full model (**Fig. S5**; Cohen’s d; Model without entropy/repetition metrics: *d* = −0.10, Model without entropy-based metrics: *d* = −0.10, Full model with entropy and repetition metrics: *d* = −0.39). The full model with entropy-based metrics was the only model that came close to replicating the magnitude of differences in deviation from matching between the 40/5 and 40/10 schedules in behavioral data (**Fig. S5c–d**).

For monkeys, predicted deviation from matching from both regression models without entropy metrics was significantly lower in the 70/30 than the 80/20 reward environment (**Fig. S5e–f**; two-sided t-test; Model without entropy/repetition metrics: *p* = 1.19 × 10^-49^, Model without entropy-based metrics: *p* = 1.32 × 10^-29^). Only the regression model with entropy metrics replicated the observed lack of difference in undermatching between reward schedules. (**Fig. S5g–h**; Full model with entropy metrics: *p* = 0.36; Observed difference between reward schedules: *p* = 0.19). Therefore, entropy-based metrics are necessary and sufficient to capture the influence of reward schedule on deviation from matching.

### Reinforcement learning models do not capture entropy-based metrics well

To capture the observed variability in entropy-based metrics and underlying learning and choice mechanisms, we next fit choice behavior using three reinforcement learning (RL) models. These models assumed different updating of reward values and one included a term to mimic the influence of previous choices on choice during the current trial (see **Methods** for more details). Out of the three studied models, we found that RL2, in which the estimated reward values for the unchosen side or stimulus decays to zero over time, provided the best fit of choice behavior for both mice and monkeys as reflected in the lowest Akaike Information Criterion (AIC) (**Fig. 5a–b**).

**Figure 5.**
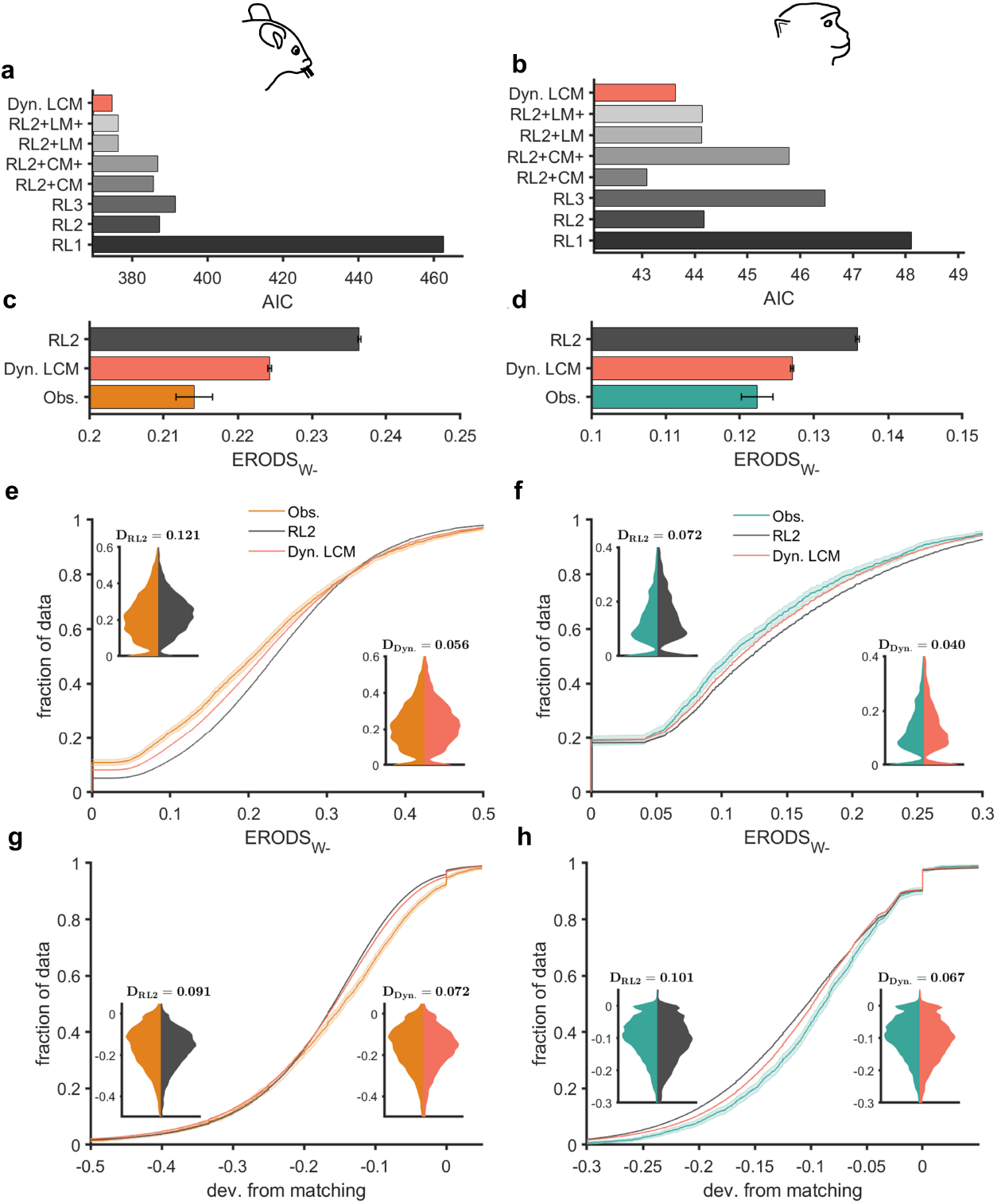
Dynamic loss-choice memory model better accounts for choice behavior and undermatching and exhibits less bias and error in capturing entropy-based metrics. **(a–b)** Comparison of goodness-of-fit of three RL models, the dynamic loss-choice memory (LCM) model, and models with some components of the dynamic LCM model. Plotted is the average Akaike Information Criterion (AIC) over all sessions for mice (a) and monkeys (b). **(c–d)** Comparison of average ERODS_W−_ observed in animals and ERODS_W−_ predicted based on simulations of the best-fitting RL model and the new dynamic LCM model. Error bars represent SEM. **(e–h)** Empirical cumulative distribution functions (CDFs) of ERODS_W−_ (e-f) and deviation from matching (g-h) observed in animals and predicted from simulations of the RL2 dynamic LCM models. Shaded bars around CDFs indicate 95% confidence interval. Insets display the distribution of observed metrics versus metrics predicted using the RL model (left inset) and dynamic LCM model (right inset). Distributions are scaled such that their maximum widths are equal. Displayed *D*-values are the test-statistic from a two-sided Kolmogorov-Smirnov test comparing the distributions. Lower D-values indicate increased similarity between distributions (less different CDFs); D-values are directly comparable because the same, large number of simulations (100 per session) were performed to estimate population distributions of metrics. The dynamic LCM model better captured deviation from matching in mice by nearly 20% and in monkeys by over 30%.

We next tested whether RL2 could replicate observed distributions of entropy-based metrics and undermatching by simulating the model during our experiment using parameters obtained from model fitting. Due to the large number of simulations performed (100 simulations per session, mice: *n* = 331,900 blocks; monkeys: *n* = 221,200 blocks), we were able to estimate population distributions of metrics for each model. We found that the average predicted ERODS_W−_ was significantly higher than the average observed ERODS_W−_, suggesting the RL model underutilizes loss-dependent and option-dependent strategies when compared to mice and monkeys in our experiments (**Fig. 5c-d)**. Additionally, block-wise predicted and observed ERODS_W−_ were both highly variable, but the distribution of predicted ERODS_W−_ was different than the observed distribution for both mice and monkeys (**Fig. 5e-f**; Two-sided Kolmogorov-Smirnov test; mice: *D* = 0.121; monkeys: *D* = 0.072). Moreover, the predicted distribution of deviation from matching was different from the observed distribution for both mice and monkeys (**Fig 5g-h**; Two-sided Kolmogorov-Smirnov test; mice: *D* = 0.091; monkeys: *D* = 0.101).

Finally, we also computed undermatching and all behavioral metrics in simulated data using RL2 with random parameter values. We found that our entropy-based metrics are better predictors for deviation from matching than the parameters of the RL2 model (see **Fig. S6**). Together, our results illustrate that RL models failed to replicate observed distribution of ERODS_W−_ and variability in matching behavior, pointing to additional mechanisms that contribute to behavior. Moreover, because ERODS_W−_ is highly correlated with undermatching in observed behavior (**Fig. 4**) and RL simulations (**Fig. S6**), our results suggest that a model that better captures ERODS_W−_ may also better capture variability in matching behavior.

### Model inspired by entropy-based metrics captures choice behavior more accurately

The deviations of predicted ERODS_W−_ from observed ERODS_W−_ suggest RL models underutilize loss-dependent and option-dependent strategies; that is, they fail to capture the influence of option (or action) and loss in the current trial on choice in the subsequent trial. To fix this problem, we developed a new model, which we refer to as the dynamic loss-choice memory (LCM) model, that augments a RL component with an option-dependent choice memory and an outcome-dependent loss memory (see **Methods**).

More specifically, the loss-memory component encourages either staying or switching in response to loss and is modulated by expected uncertainty. Here, expected uncertainty is defined as the expected unsigned reward prediction error in the RL component. As uncertainty increases, the weight of the loss-memory component increases proportionately, making choice more dependent on previous reward feedback. In contrast to RL models with adaptive learning rates (e.g., as in Pearce-Hall model), changes in the loss-memory component only influence subsequent choice, not the aggregation of values in the RL component of our model. The choice-memory component encourages either staying on or switching from options that have been chosen recently. Since the model typically chooses the option with a higher value, the choice-memory component captures strategy in response to selection of the better or worse option. Importantly, our model is agnostic about the strategy employed by the loss- and choice-memory components; for example, the loss-memory component could encourage lose-stay or lose-switch behavior, and the choice-memory component could encourage stay or switch after selection of the worse option (or action), which we refer to as worse-stay and worse-switch. In summary, the loss-memory component can capture lose-switch strategies and the choice-memory component can capture worse-switch strategies. The addition of these components to an RL component creates a model that can flexibly modulate reward- and option-dependent strategies, which may increase the consistency of strategy in response to loss on the worse option (reduce ERODS_W−_).

We found that the dynamic loss-choice memory model fit choice behavior better than all previous RL models as indicated by a lower AIC (**Fig. 5a-b**). Also, the dynamic LCM model explained a large absolute proportion of variance in choice behavior as indicated by McFadden pseudo R^2^ values of 0.35 for mice and 0.72 for monkeys (**Table S2**). As expected, the dynamic loss-choice memory model better captured the average consistency of strategy in response to loss on the worse option or action (**Fig. 5c-d**). Importantly, the new model also better captured the observed distribution of ERODS_W−_ in both mice and monkeys (**Fig 5e-f**; two-sided Kolmogorov-Smirnov test; mice: *D* = 0.056; monkeys: *D* = 0.040). This improvement in capturing ERODS_W−_ corresponded with similar improvements in capturing deviation from matching. The predicted distribution of deviation from matching from the dynamic LCM model better replicated the observed distribution of deviation from matching than the predicted distribution from the best-fitting RL model in both mice and monkeys (**Fig 5g-h**; Two-sided Kolmogorov-Smirnov test; mice: *D* = 0.072; monkeys: *D* = 0.067). This improvement is significant; there is a nearly 20% reduction in the maximum difference between cumulative distribution functions (CDFs) for mice and an over 30% reduction in the maximum difference between CDFs for monkeys.

To identify the contributions of each component of the dynamic LCM model in capturing choice behavior, we also tested four additional models that are composed of RL2 and either the loss-memory or the choice-memory components of the dynamic LCM model. In two of these models, RL2+LM+ and RL2+CM+, the weights of the loss-memory and choice-memory components were restricted to positive values. We found that in mice, the dynamic LCM model fit trial-by-trial choice behavior and captured global undermatching better than models that include either of the two components (**Fig. 5a; Table S2**). Moreover, improvement in capturing local and global choice behaviors in mice was larger when including loss-memory component than choice-memory component, pointing to a larger contribution of loss-dependent strategies in mice. In monkeys, however, goodness-of-fit using the RL2+CM model was indistinguishable from goodness-of-fit based on the dynamic LCM model, suggesting the importance of option-dependent strategy for explaining choice behavior in monkeys (**Fig. 5a; Table S2**). The fit of choice behavior using choice-memory component with positive values only (RL2+CM+ model), was worse than that of the simple RL2 model, whereas bidirectional choice-memory component improved the fit of both local and global choice behaviors (**Fig 5a; Table S2**). These results indicate that bidirectional weighting inspired by entropy-based metrics was responsible for most of the observed improvements in capturing choice behavior in monkeys.

Finally, to understand how the dynamic LCM model modulates specific loss- and option-dependent strategies in mice and monkeys, we compared the distributions of model parameters between the dynamic LCM, RL2, RL2+CM, and RL2+LM models. Interestingly, the loss-memory and choice-memory components both had positive average weights in mice and negative average weights in monkeys (**Fig S7i-l)**. In other words, the loss-memory component generally resulted in lose-switch strategies in mice and lose-stay strategies in monkeys, whereas the choice-memory component encouraged worse-switch strategies in mice and worse-stay strategies in monkeys. We also found that in mice, the average weights of the choice- and reward-memory components were higher in the dynamic LCM model than in the RL2+CM and RL2+LM models, respectively, indicating that the two components may interact to modulate behavior (**Fig S7i-j)**. This dovetails with our entropy-based results that show response to the loss after selection of the worse option is uniquely important for global choice behaviors.

To summarize, the proposed model improved fit of choice behavior and captured our metrics and undermatching more accurately, whereby revealing that undermatching behavior arises from competition among multiple components incorporating loss-memory and choice-memory on a trial-by-trial basis. Importantly, we used deviations in predicted entropy-based metrics from their observed values to identify shortcomings in existing RL models and propose mechanisms to mitigate them.

## Discussion

Undermatching is a universal behavioral phenomenon that has been observed across many species. Here, we show that novel entropy-based metrics based on response to reward feedback can accurately predict undermatching in mice and monkeys, suggesting that inconsistent use of local reward-dependent and option-dependent strategies can account for a large proportion of variance in global undermatching. Moreover, we demonstrate that these entropy-based metrics can be utilized to construct more complex reinforcement learning models that are able to capture choice behavior, undermatching, and utilization of reward-dependent strategy. Together, our entropy-based metrics provide a powerful model-free tool to develop and refine computational models of choice behavior and reveal neural mechanisms underlying adaptive behavior.

Similar to many previous studies of matching behavior (Herrnstein and Loveland, 1974; William, 1979; Belke and Belliveau, 2001; Sugrue et al., 2004; Tsutsui et al., 2016), we observed significant, but highly variable undermatching in both mice and monkeys. Although many studies (William, 1979; Otto et al., 2011; Iigaya et al., 2019) have speculated a relationship between local choice and learning strategies and global matching behavior, few aimed to predict observed undermatching (except Iigaya et al., 2019). By focusing on the variability in undermatching, here, we were able to show that global undermatching can be largely explained by the degree of inconsistency in response to no reward on the worse option (ERODSw-) across species. Specifically, ERODSw- could explain about 50% and 40% of the variance in undermatching in mice and monkeys, respectively. The proposed entropy metrics were able to predict undermatching across two very different species despite differences in the tasks including utilized reward probabilities and schedules (40/5 and 40/10 probabilities with baiting vs. complementary 80/20 and 70/30 with no baiting), learning modality (action-based vs. stimulus-based), choice readout (licks vs. saccades), and predictability of block switches (unpredictable vs. semipredictable), suggesting the proposed metrics are powerful and generalizable.

The proposed entropy-based metrics complement and improve upon commonly used behavioral metrics such as win-stay, lose-switch, and the U-measure (Miller and Frick, 1949). Although the U-value has been used to measure consistency or variability in choice behavior (Machado, 1992, 1993), it is difficult to interpret and fails to capture sequential dependencies in choice (Kong et al., 2017). Proposed entropy-based metrics avoid these issues because they have both clear interpretations and can capture the sequential dependence of choice on previous reward and/or selected action or option. Similarly, although win-stay and lose-switch provide valuable information (Bari et al., 2010; Otto et al., 2011; Dalton et al., 2014, 2016; Jang et al., 2015; Gruber and Thapa, 2016), these probabilities do not solely reflect the effects of reward feedback on staying (or similarly switching) as they both depend on the probability of stay. For example, if staying behavior is independent of reward, win-stay and lose-switch values simply reflect the overall stay and switch probabilities, respectively. We also found that win-stay and lose-switch are not strong predictors of undermatching because their relative importance depends on the overall probability of winning. For example, if the overall probability of reward is high, lose-switch is less useful for predicting behavior because response to loss represents strategy in a small subset of trials. Therefore, win-stay and lose-switch cannot capture the degree to which staying and switching strategy depend on reward outcome only. The entropy-based metrics such as ERDS overcome these issues by combing win-stay and lose-switch with p(win) and p(stay). Finally, more complex model-independent metrics involving conditional probabilities (Ito and Doya, 2009) are not desirable because the number of possible conditional probabilities grows exponentially with the number of conditioning variables (e.g., reward, side, stimulus) and the number of preceding trials examined. Entropy-based metrics can overcome this issue by capturing sequential information in a single quantity because additional entropy-based metrics can easily be defined based on longer sequences of choice, reward, other task variables.

As shown by multiple studies, models that fit choice data best may still fail to replicate important aspects of behavior (Palminteri et al., 2017; Wilson and Collins, 2019). Therefore, model validation must involve analyzing both a model’s predictive potential (fitting) and its generative power (replication of behavior in simulations). We used shortcomings of existing RL models in capturing the most predictive entropy-based metrics to detect additional mechanisms underlying adaptive behavior. This approach can be applied to other tasks in which similar or different entropy-based metrics are most predictive of global choice behavior (matching or other metrics). Our aim here was not to find the best model for capturing all aspects of behavior but instead to provide a framework for how response to reinforcement can be used both to explain global behavior and to capture local choice behavior.

Using this method, we constructed a new dynamic loss-choice memory model that augments a reinforcement-learning system with a choice-memory component that captures option-dependent strategies and a loss-memory component that captures loss-dependent strategies. Previous studies have also shown that a combination of win-stay lose-switch (WSLS) strategies with RL models could improve fit of choice behavior and the average matching behavior (Ito and Doya, 2009; Otto et al., 2011; Worthy and Maddox, 2014). The choice-memory component used here is similar to choice kernel models that have been shown to improve fit of choice behavior (Lau and Glimcher, 2005; Wittmann et al., 2020). Nonetheless, the proposed dynamic loss-choice memory model is a novel combination of these components. Critically, the weights of the loss and choice components could be either positive or negative. This parallels how entropy-based metrics capture response to reward feedback considering that low entropy can result from strong positive or negative influences of recent rewards or choices (e.g., high win-stay and high win-switch both correspond to low entropy). Because of this mechanism, the model can explain a few findings not related to our study. For instance, the model can explain how variability in choice can be enhanced (Machado, 1992) by assigning a negative weight to the choice-memory system that encourages switching from recently selected options. Moreover, the model can explain why high levels of global reward can decrease lose-switch (Wittmann et al., 2020): a negative weight in the reward-memory component based on expected unsigned reward prediction error results in increased influence of the component in unrewarded trials (decreasing lose-switch) when global reward rate is high. The model may also facilitate quick adaptation to reversals, a behavior that has previously been explained using Bayesian approaches (Costa et al., 2015), because negative weights in the choice-memory component and in the reward-memory component weighted by unsigned reward prediction error both encourage faster response to reversals. Neural correlates of a similar loss-memory component weighted by recent reward prediction errors have been identified in the dorsal anterior cingulate cortex of humans (Wittmann et al., 2016). Moreover, neural correlates of such choice memories have been identified in various cortical areas of monkeys including the dorsolateral prefrontal cortex, dorsal medial prefrontal cortex, lateral intraparietal area, and the anterior cingulate cortex (Barraclough et al., 2004; Seo et al., 2009; Spitmaan et al., 2020).

In summary, we show that entropy-based metrics are good predictors of global choice behavior across species and can be used to refine reinforcement learning models. Results from fitting and simulating the dynamic loss-choice memory model suggest that recent choices and rewards affect decisions in ways beyond their influence on the update of the stimulus or action values. Thus, entropy-based metrics have the potential to open a new realm of possibilities for understanding neural mechanisms underlying adaptive behavior.

## Methods

### Experimental paradigm in mice

Mice performed a dynamic foraging task in which after receiving a “go cue” signaled by an odor, they licked one of the two water tubes (on left and right) to harvest possible reward. In 5% of trials a “no-go cue” was presented by another odor signaling that licks would not be rewarded or punished. If a mouse licked one of the tubes after a go-cue odor, reward was delivered probabilistically. Each trial was followed by an inter-trial interval drawn from an exponential distribution with a rate parameter of 0.3.

The reward probabilities assigned to the left and right tubes were constant for a fixed number of trials (blocks) and changed throughout the session (block switches). Block lengths were drawn from a uniform distribution that spanned a range of 40-100 trials, however, the exact block lengths spanned smaller ranges for individual sessions, resulting in variable block lengths with most block lengths ranging between 40-80 trials. If mice exhibited strong side-specific biases, block lengths were occasionally shortened or lengthened. “Miss trials,” where the mouse did not make a choice, were excluded for all analyses described here.

Mice performed two versions of the task, one with 16 different sets of reward schedules and another with two sets of reward schedules. The vast majority (469 out of 528) of sessions used two sets of reward probabilities equal to 0.4 and 0.1, and 0.4 and 0.05, which we refer to as 40/10 and 40/5 reward schedules. Here, we focus on the most frequent blocks (40/5 and 40/10 reward schedules). Rewards were “baited” such that if reward was assigned on a given side and that side was not selected, reward would remain on that side until the next time that side was selected. Due to this baiting mechanism, the probability of obtaining reward on the unchosen side increased over time as during foraging in a natural environment. Therefore, optimal performance in the task generally required a strategy that involved selecting options in proportion to their reinforcement rates (perfect “matching” behavior). In total, 16 mice performed 469 sessions of the two-probability version of the task for a total of 3319 blocks (1786 and 1533 blocks with 40/5 and 40/10 reward schedules, respectively) and 189,199 trials. More details about the experimental set up are provided in Bari et al (Bari et al., 2019).

### Experimental paradigm in monkeys

In the reversal learning task in monkeys, Costa and colleagues (Costa et al., 2016) trained monkeys to fixate on a central point on a screen to initiate each trial (**Fig. 1**). After fixation, two stimuli, a square and circle, were presented on the screen to the left and right of the fixation point. The side that the stimuli were presented on was chosen randomly and was not related to reward. Monkeys made saccades to a stimulus and fixated on the stimulus to indicate their choice in each trial. A juice reward was delivered probabilistically via a pressurized tube based on the chosen stimulus. Each trial was followed by a fixed 1.5s inter-trial interval. Trials in which the monkey did not make a choice or failed to fixate were immediately repeated.

Monkeys completed sessions that contained around 1300 trials on average divided into superblocks of 80-trials. Within each superblock the reward probabilities assigned to each cue were reversed randomly between trials 30 and 50, such that the stimuli that was less rewarding at the beginning of the superblock became more rewarding and vice versa. Every 80 trials, monkeys were presented with new stimuli that varied in color but not shape. Monkeys performed two variants of the task, a stochastic variant with three reward schedules (80/20, 70/30, and 60/40) and a deterministic variant with one reward schedule (100/0). Here, we focus our analyses on the 80/20 and 70/30 reward schedules (2212 blocks of the task performed by 4 monkeys) as they provide two levels of uncertainty similar to the experiment in mice. For more details, please see Costa et al. (Costa et al., 2015, 2016).

### Behavioral metrics

#### Matching Performance

To measure the overall response to reinforcement on the two choice options (e.g., left and right actions when reward is based on the location) in each block of the experiment, we defined undermatching (UM) as:

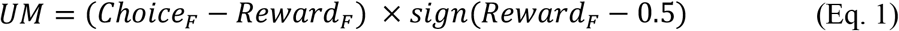

where *sign*(0) = 1 and choice and reward fractions (*Choice_F_, Reward_F_*) are defined as follows:

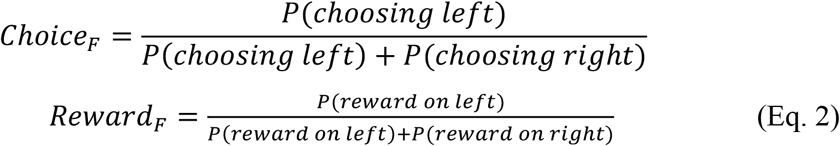

Therefore, UM measures the difference between choice and reward fractions toward the more rewarding options (side or stimulus). Similarly, UM can be computed based on the color of stimuli when color is informative about reward outcome. Based on our definition, negative and positive values for UM correspond to undermatching and overmatching, respectively.

#### Win-stay and lose-switch

Win-stay (WS) and lose-switch (LS) measure the tendency to repeat a rewarded choice (in terms of action or stimulus) and switch away from an unrewarded choice, respectively. These quantities are based on the conditional probabilities of stay and switch after reward and no reward, respectively, and can be calculated in a block of trials as follows:

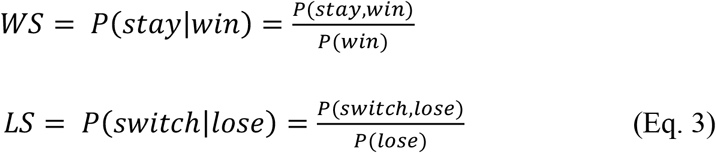

where *P*(*win*) and *P*(*stay*) are the probabilities of harvesting reward and choosing the same side (or stimulus) in successive trials, *P*(*lose*) = 1 — *P*(*win*), and *P*(*switch*) = 1 — *P*(*stay*).

#### Repetition Index

Repetition index (RI) measures the tendency to repeat a choice beyond what is expected by chance and can be computed by subtracting the probability of stay by chance from the original probability of stay (Soltani et al., 2013). RI can be computed based on the repetition of left or right choices as follows:

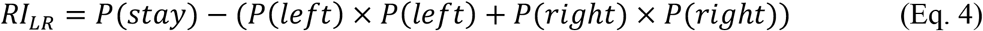

In general, RI reflects a combination of reward-dependent and reward-independent strategies as well as the stochasticity in choice (temperature in the logit function translating value difference to choice probability).

Repetition index can also be measured based on other option or choice attributes that predict reward such as the color of the chosen option. For example, RI can be defined based on selection of the better and worse options (*RI_BW_*) when such options exist in a task:

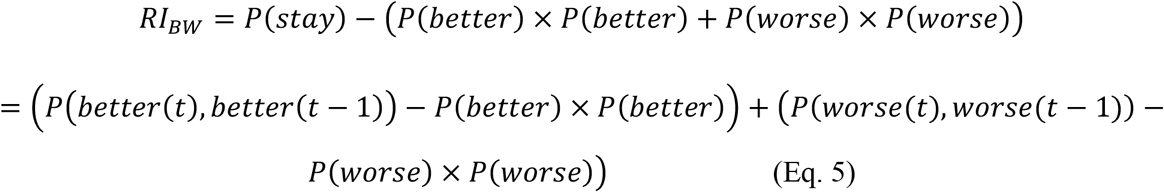

where *t* is the trial number. Using Eq. 5, *RI_BW_* can be decomposed into two pieces, *RI_B_* and *RI_W_*, that measure the tendency to repeat the better and worse option, respectively:

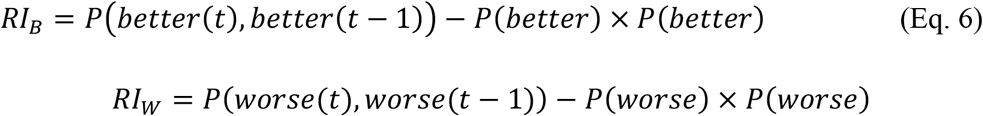

#### New entropy-based measures

In order to quantify the influence of previous reward outcome on choice behavior in terms of stay or switch, we defined the conditional entropy of reward-dependent strategies (ERDS) that combine tendencies of win-stay and lose-switch into a single metric. More specifically, ERDS is defined as the conditional entropy of using stay or switch strategy depending on win or lose in the preceding trial:

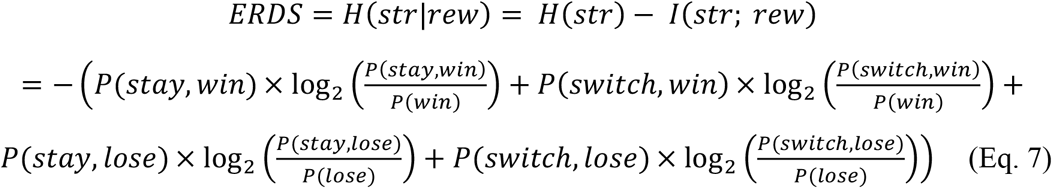

where *H*(*str*) and *I* (*str; rew*) are entropy of strategy (stay or switch) and mutual information of strategy and reward:

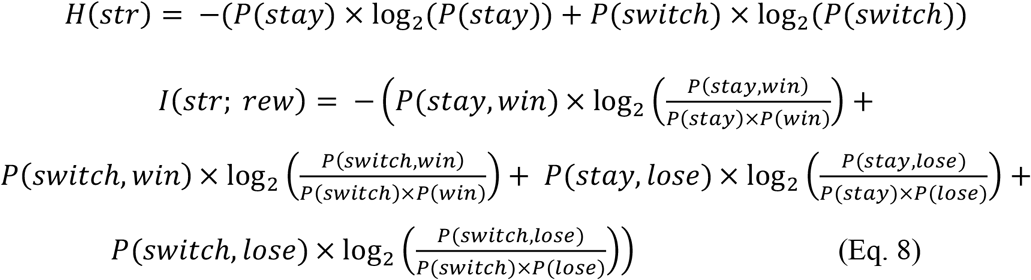

To better show the link between ERDS and win-stay, lose-switch, and *p*(*win*), Eq. 7 can be rewritten as follows:

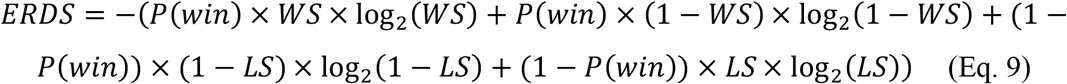

ERDS can be decomposed into two components, *ERDS_+_* and *ERDS_−_* (*ERDS* = *ERDS_+_* + *ERDS_−_*), to allow separation of animals’ response to rewarded (win) and unrewarded (loss) outcomes:

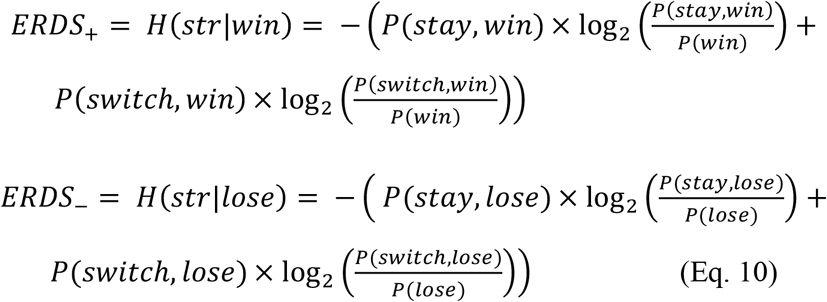

The above equations also show that *ERDS_+_* and *ERDS_−_* are linked to win-stay and lose-switch, respectively.

Considering that RI can be decomposed to repetition after better and worse options (Eq. 5), and following the same logic used to derive ERDS, one can define the conditional entropy of option-dependent strategy (EODS) based on staying on or switching from the better or worse option (or left and right option) in two consecutive trials:

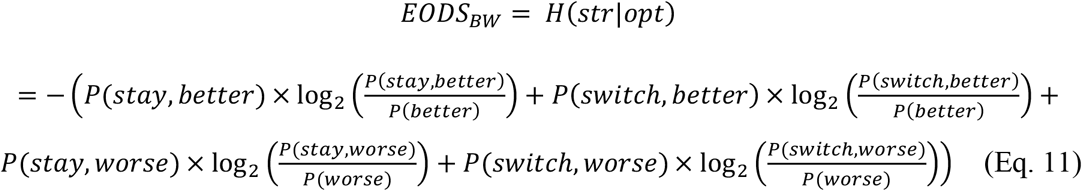

*EODS_BW_* can be decomposed into two components based on the better and worse options:

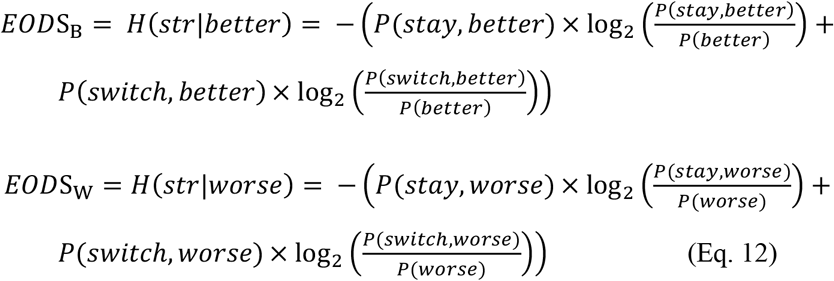

EODS can also be computed based on other choice attributes (e.g., location or color of options) to examine what information drives stay or switch behavior:

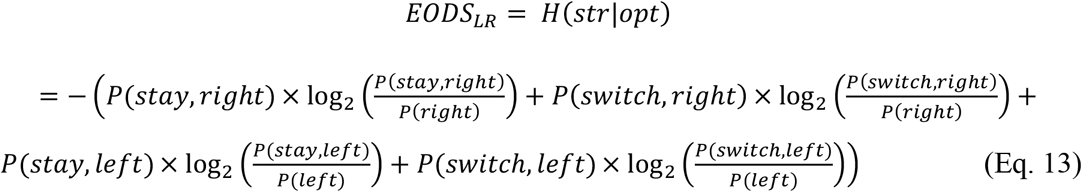

We should note that ERDS and EODS are directly comparable and provide insight into the consistency of strategy adopted by an animal. Lower ERDS than EODS suggests that an animal’s decisions are more consistently influenced by immediate reward feedback than selection of the better or worse option. A lower EODS than ERDS suggests the opposite. It is worth noting that ERDS decompositions (ERDS+ or ERDS-) cannot be directly compared to EODS decompositions (EODS_B_ or EODS_W_) because they encompass different sets of trials; that is, trials where the animal wins may not be trials where the animal chooses the better side and vice versa.

Because conditional entropies can be defined for any two discrete random variables, ERDS and EODS can be generalized to combinations or sequences of combinations of reward and option. Hence, we can define the entropy of reward- and option-dependent strategy (ERODS), a measure of the dependence of strategy on option and reward.

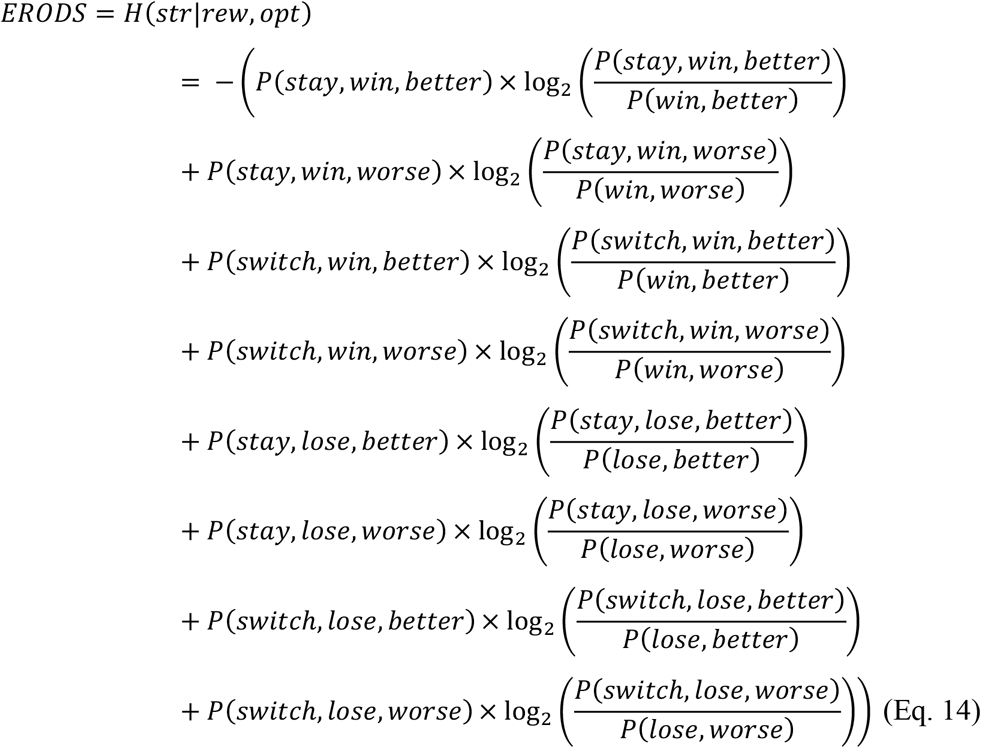

ERODS can be decomposed based on choosing the better or worse option in the previous trial, winning or losing in the previous trial, or combinations of option and reward (e.g., choose better option and win on the previous trial).

Decomposing ERODS based on reward option combinations gives:

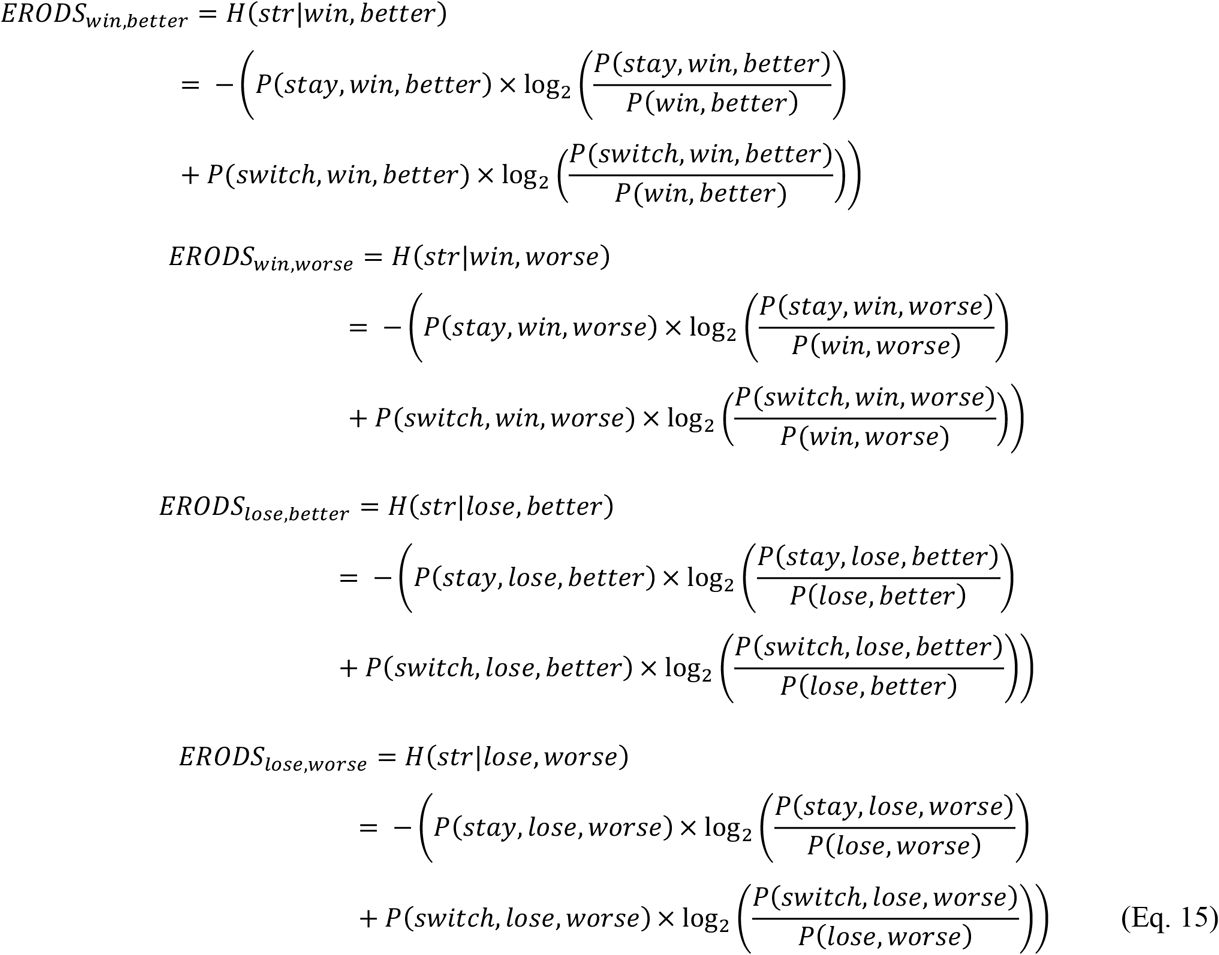

Finally, ERODS can also be decomposed based on selection of the better or worse option:

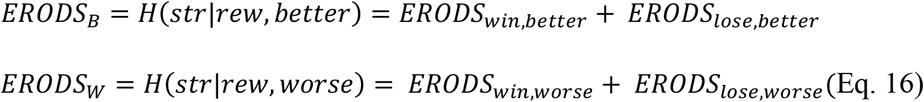

or winning or losing in the previous trial:

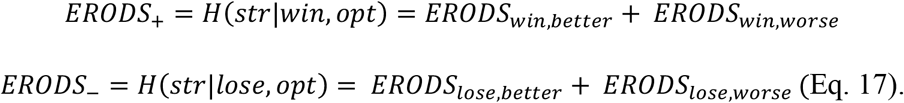

### Reinforcement learning models

We used three reinforcement learning models to fit choice behavior. In all these models, reward probabilities or values associated with the right and left sides (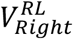 and 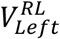) for mice or circle and square stimuli (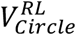 and 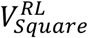) for monkeys were updated differently depending on whether a given choice was rewarded or not. In the first model, which we refer to as RL1, only the estimated reward probability associated with the chosen side or stimulus 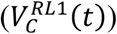 was updated as follows:

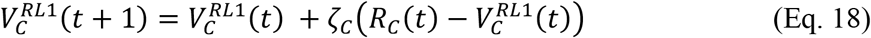

where *C* ∈ {*Left, Ripht*} for mice and *C* ∈ {*Circle, Square*} for monkeys, *R_C_*(*t*) =1 or 0 indicates reward outcome on trial *t*, and *ζ_C_* corresponds to the learning rate (*α_rew_ and α_unrew_*) depending on the whether the choice was rewarded or not rewarded.

In the second model (RL2), the estimated reward probability associated with the unchosen side or stimulus 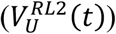 was also updated as follows:

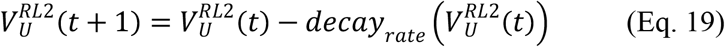

where *decay_rate_* is the decay (or discount) rate of value of the unchosen side or stimulus, corresponding to forgetting reward values over time. The third model (RL3) was similar to the first model in terms of the integration of reward outcomes and updating of estimated reward probabilities but had an additional choice-memory component for capturing the effects of previous choices. More specifically, the choice-memory value for the chosen side or stimulus in trial *t*, 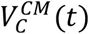, and for the unchosen side or stimulus in trial *t*, 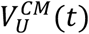, are updated as follows:

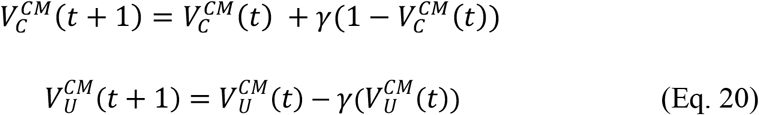

where *γ* represents the decay rate for the choice-memory value. In RL3, the overall reward value for the chosen (unchosen) side/stimulus on trial *t, V_C_*(*t*) (*V_U_* (*t*)), is a linear sum of the choice-memory value and the estimated reward probability for RL1:

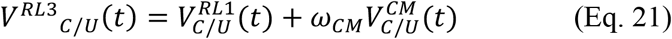

where *ω_CM_* determines the relative contribution of the choice-memory component and 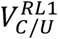 is the estimated reward probability of the chosen/unchosen side or stimulus based on RL1.

For all three models, the probability of selecting the left (or right) side is represented as a sigmoid function of the difference in estimated reward probabilities or overall reward values for the left and right sides. Hence, the estimated probability of choosing the left side for mice (or circle for monkeys) in trial *t, P_*Left*(*circle*)_*(*t*), is:

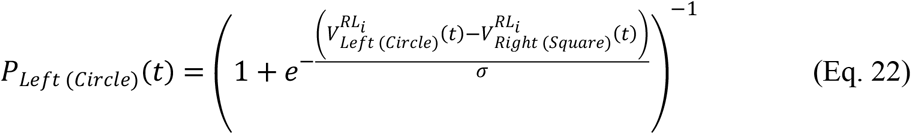

where *σ* quantifies the stochasticity of choice and *RL_i_* is RL1, RL2, or RL3. Values of *decay_rate_, γ, α_rew_*, and *α_unrew_* ranged from 0 to 1 for all three models, and values of *σ* ranged from 0 to 100. These models were selected because previous research has suggested they can replicate matching or undermatching phenomena (Sugrue et al., 2004; Soltani and Wang, 2006).

As we show here, however, these models did not replicate important behavioral phenomena in our data (see **Results**). Therefore, we developed a new model based on entropy-based metrics. The new model augments RL2 with a loss-memory component to capture loss-dependent strategy and a choice-memory component to capture option-dependent strategy. In the choice-memory component, the choice-memory value of the chosen (unchosen) option (*V^CM^ _C/U_*) was updated as in RL3, with *γ* = mean(*α_rew_, α_unrew_*) as the decay rate of the choice-memory component.

In the loss-memory component, the value of the chosen side is the expected reward prediction error on the next trial in unrewarded trials and 0 in rewarded trials. Hence, the expected reward prediction error (*E_rpe_*) and the loss-memory value of the chosen side (*V^LM^_C_(*t*)*) are updated as follows:

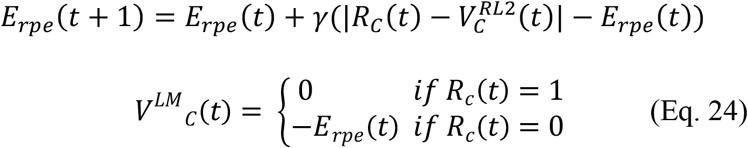

where *V^RL2^_C_*(*t*) is the value of the chosen side in RL2. In the full model (dynamic loss-choice memory, LCM), the overall reward values of the chosen and unchosen sides in trial *t, V_C_* (*t*) and *V_U_*(*t*), are computed as follows:

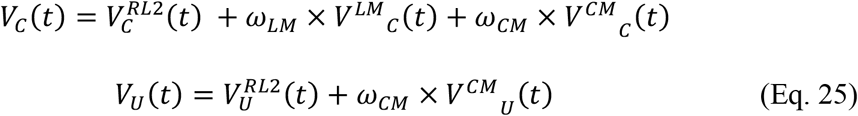

where *ω_LM_* and *ω_CM_* are free parameters that determine the relative weight of the reward-memory and choice-memory component, respectively. Values of *ω_LM_* and *ω_CM_* varied from −1 to 1, such that the effects of recent reward and choice on future choice could increase either staying or switching behavior. Notably, *V^RL2^_C/U_*(*t*) are independent of *V^LM^_C_*(*t*) and *V^CM^_C_*(*t*), such that the values in the reward-memory and choice-memory components only influence choice in the subsequent trial. Finally, the probability of choosing a side for mice (or a stimulus for monkeys) is computed using a sigmoid function of difference in the overall reward values as described above.

To test the components of the full model individually and the effects of allowing negative parameter weights, we fitted four additional control models. The RL2+LM model is the full model without the choice-memory component, and the RL2+CM model is the full model without the loss-memory component. In the RL2+LM+ and RL2+CM+ models, however, *ω_LM_* and *ω_CM_* range from 0 to 1 instead of from −1 to 1.

### Model fitting and simulations

We used the standard maximum likelihood estimation method to fit and estimate the best-fit parameters for the models described above. To quantify goodness-of-fit, we computed the Akaike Information Criterion (AIC):

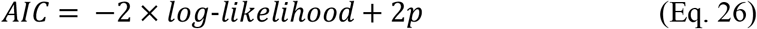

where *p* is the number of free parameters in a given model.

To quantify an absolute measure of goodness of fit, we computed the McFadden R^2^ (McFadden, 1973) for each model:

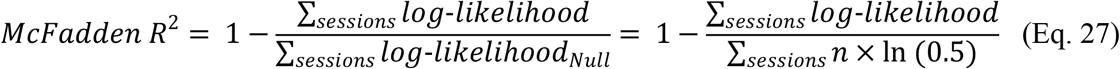

where *n* is the number of trials in a given session. One hundred model simulations were performed per session using best-fit parameters. The large number of simulations allowed us to estimate the population distributions of all metrics. Finally, we conducted additional simulations of RL2 using random parameter values to examine the relationship between parameters and entropy-based metrics. For these simulations, *α_rew_, α_unrew_*, and *σ* varied in the range of (0, 1) and *decay_rate_* was set to 0.1.

### Data analyses and stepwise regressions

Stepwise regressions were conducted using MATLAB’s stepwiselm and stepwisefit functions. The criterion for adding or removing terms from the model was based on an F-test of the difference in sum of squared error resulting from the addition or removal of a term from the model. A predictor was added to the model if the *p*-value of the F-test was less than 0.0001, and a predictor was removed from the model if the *p*-value of the F-test was greater than 0.00011. The order in which predictors were added to the stepwise regression models was as follows.

#### Model without entropy/repetition metrics (mice)

In the first step of the stepwise regression, *p*(*win*) entered the regression equation as a significant predictor of undermatching, *RMSE* = 0.1119,*p* = 4.12 × 10^-175^. In the next steps, *p*(*stay*) (*RMSE* = 0.1053,*p* = 2.41 × 10^-89^), and *p*(*stay|win*) (*RMSE* = 0.1046, *p* = 7.41 × 10^-11^) entered the regression equation.

#### Model without entropy/repetition metrics (monkeys)

In the first step of the stepwise regression, *p*(*stay*) entered the regression equation as a significant predictor of undermatching, *RMSE* = 0.0599,*p* = 1.31 × 10^-160^. In the next steps, *p*(*win*) (*RMSE* = 0.0579,*p* = 1.86 × 10^-35^), *p*(*switch|lose*) (*RMSE* = 0.0575,*p* = 2.85 × 10^-8^), and *p*(*stay|win*) (*RMSE* = 0.0571,*p* = 5.61 × 10^-9^) entered the regression equation. In the final step, *p*(*stay*) was removed from the equation because it was no longer a significant predictor (*RMSE* = 0.0572,*p* = 6.37 × 10^-4^).

#### Model without entropy metrics (mice)

In the first step of the regression process, *p*(*win*) entered the model (*RMSE* = 0.1119,*p* < 4.12 × 10^-175^). In the next steps, *p*(*stay*)(*RMSE* = 0.1053,*p* < 2.41×10^-89^),RI_W_ (*RMSE* = 0.0929,*p* < 4.80 × 10^-181^),*p*(*stay|win*)(*RMSE* = 0.0914,*p* < 1.03 × 10^-24^), and *RI_B_*(*RMSE* = 0.0910, *p* < 2.98 × 10^-8^) were added to the regression equation.

#### Model without entropy metrics (monkeys)

In the first step of the regression process, *p*(*stay*) entered the model (*RMSE* = 0.0599,*p* = 1.31 × 10^-160^). In the next steps, *RI_B_*(*RMSE* = 0.0505,*p* = 6.48 × 10^-167^) and *p*(*switch|lose*)(*RMSE* = 0.0503,*p* < 8.59 × 10^-5^) were added to the regression equation.

#### Full model (mice)

In the first step of the regression process *ERODS_W−_*, entered the regression equation as a significant predictor of deviation from matching, *RMSE* = 0.0717, *p* < 10^-300^. Next, entered the regression equation, *RMSE* = 0.0681, *p* < 1.61 × 10^-65^. In the following steps, EPODS_B+_ (*RMSE* = 0.0666, *p* < 1.97 × 10^-29^),*ERDS*_+_ (*RMSE* = 0.0658,*p* < 4.11 × 10^-18^), (*RMSE* = 0.0651,*p* < 1.72 × 10^-15^), *P*(*switch|lose*) (*RMSE* = 0.0639, *p* < 7.54 × 10^-24^), (*RMSE* = 0.0617,*p* < 1.46 × 10^-46^), *P*(*stay|win*)(*RMSE* = 0.0585,*p* < 1.81 × 10^-67^), *ERDS_−_* (*RMSE* = 0.0573, *p* < 5.15 × 10^-28^), *ERDS_* (*RMSE* = 0.0568,*p* < 9.59 × 10^-13^), *p*(*win*) (*RMSE* = 0.0566,*p* < 3.75 × 10^-6^), and *EODS_B_*(*RMSE* = 0.0564, *p* < 3.28 × 10^-5^) were added to the regression equation. In the final step, *EPODS_B+_* (*RMSE* = 0.0565, *p* < 5.32 × 10^-3^) was removed from the equation.

#### Full model (monkeys)

In the first step of the regression process, *EPODS_W__* entered the regression equation as a significant predictor of deviation from matching, *RMSE* = 0.0589, *p* < 7.31 × 10^-114^. Next, entered the regression equation, *RMSE* = 0.0551,*p* < 1.50 × 10^-41^. In the following steps, *EPODS_B+_* (*RMSE* = 0.0525, *p* < 2.15 × 10^-31^), *RI_B_* (*RMSE* = 0.0512,*p* < 1.09 × 10^-16^), *P*(*switch|lose*)(*RMSE* = 0.0477,*p* < 1.10 × 10^-43^), *P*(*stay|win*) (*RMSE* = 0.0468,*p* < 2.02 × 10^-13^), and *ERDS_+_* (*RMSE* = 0.0463, *p* < 5.44 × 10^-8^) were added to the regression equation.

We note that there were fewer blocks used in the full model stepwise regression because some of the specific entropy metrics were not defined for certain blocks, e.g., if a mouse or monkey never won on the worse side in a block, then ERODS_W_+ was undefined for that block. This resulted in the exclusion of around 500 blocks for mice and 700 blocks for monkeys in the final regression.

We also conducted ten-fold cross-validated regressions to predict deviation from matching (**Fig. S5**) using MATLAB’s fitrlinear and kfoldPredict functions. More specifically, stepwise regressions were performed on a set of possible predictors to determine which predictors to include in the final regression model. Then, cross-validated regressions were computed to predict deviation from matching using the set of predictors included in the final stepwise regression model.

## Acknowledgments

This work is supported by the National Institutes of Health (Grants R01DA047870 to A.S. and R01NS104834 to J.Y.C.).

---

**Figure S1.**
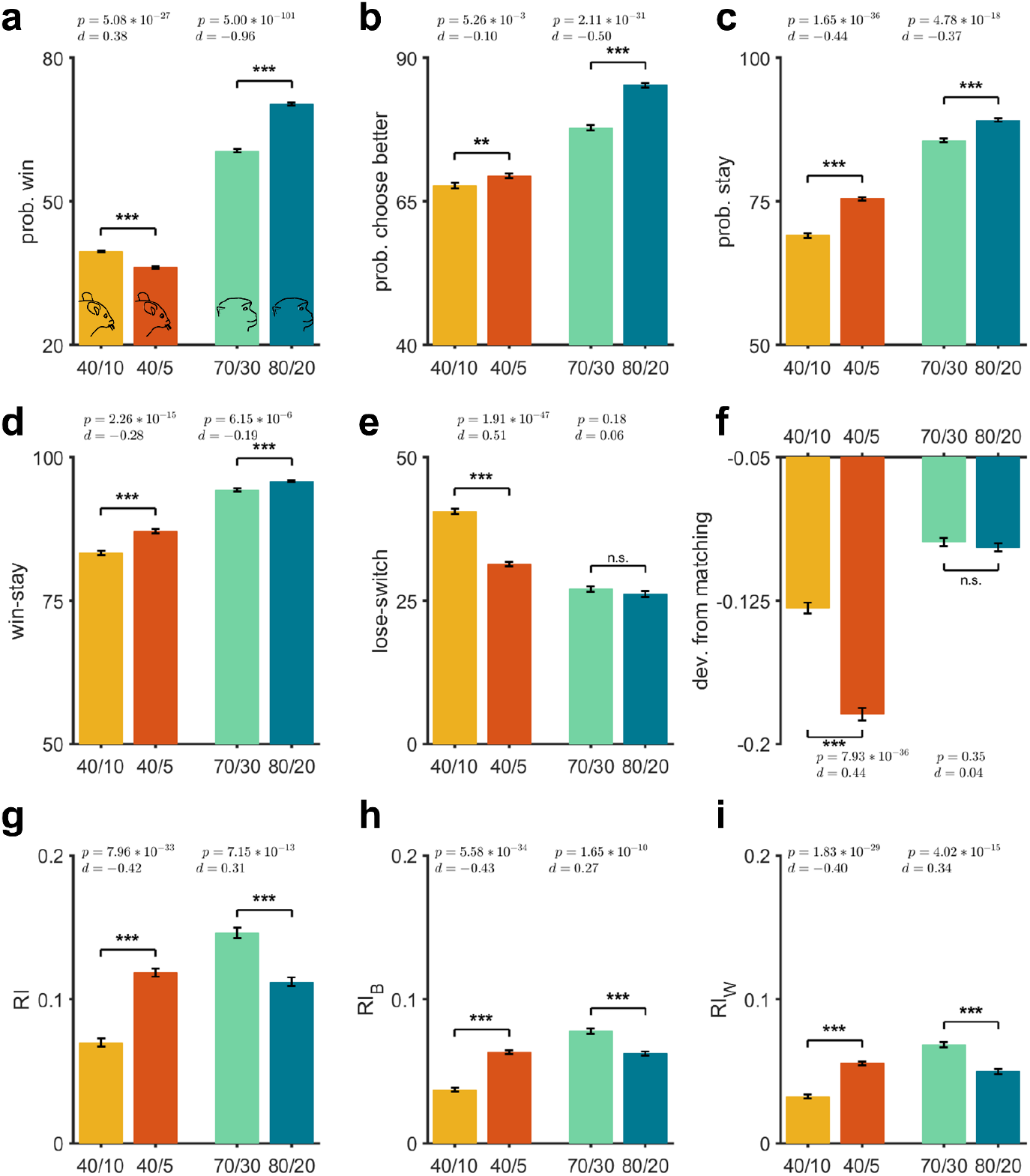
Undermatching and behavioral metrics depend on reward probabilities. Plotted are the probability of winning (a), the probability of choosing the better option (b), probability of staying on previous choice (c), win-stay (d), lose-switch (e), a measure of undermatching (f), and RI and RI decompositions (g-h) in the 40/10 and 40/5 reward schedules for mice and the 70/30 and 80/20 reward schedules for monkeys. Error bars indicate s.e.m. For mice, *n*_40/3_ = 1786 blocks and *n*_40/10_ = 1533 blocks, and for monkeys, *n*_70/30_ = 1110 blocks and *n*_80/20_ = 1102 blocks. The asterisk indicates a significant difference between the two environments using two-sided t-test with *p*-value and Cohen’s *d*-value reported on each panel. In mice, the probability of winning was significantly higher in the 40/10 schedule despite a lower probability of choosing the better side. In monkeys, the probability of winning and probability of choosing better was higher in the 80/20 schedule. Moreover, the probability of staying and the repetition index were both significantly lower in the 40/10 schedule in mice and 70/30 schedule in monkeys because the reward probabilities for the two options are more similar. Finally, both win-stay and lose-switch were closer to 0.5 in the 40/10 schedule for mice, and win-stay was closer to 0.5 in the 70/30 schedule for monkeys which may indicate a decrease in the dependence of staying and switching behavior on reward. However, the differences in *p*(*win*) and p(stay) between these environments make interpreting win-stay and lose-switch in isolation challenging.

**Figure S2.**
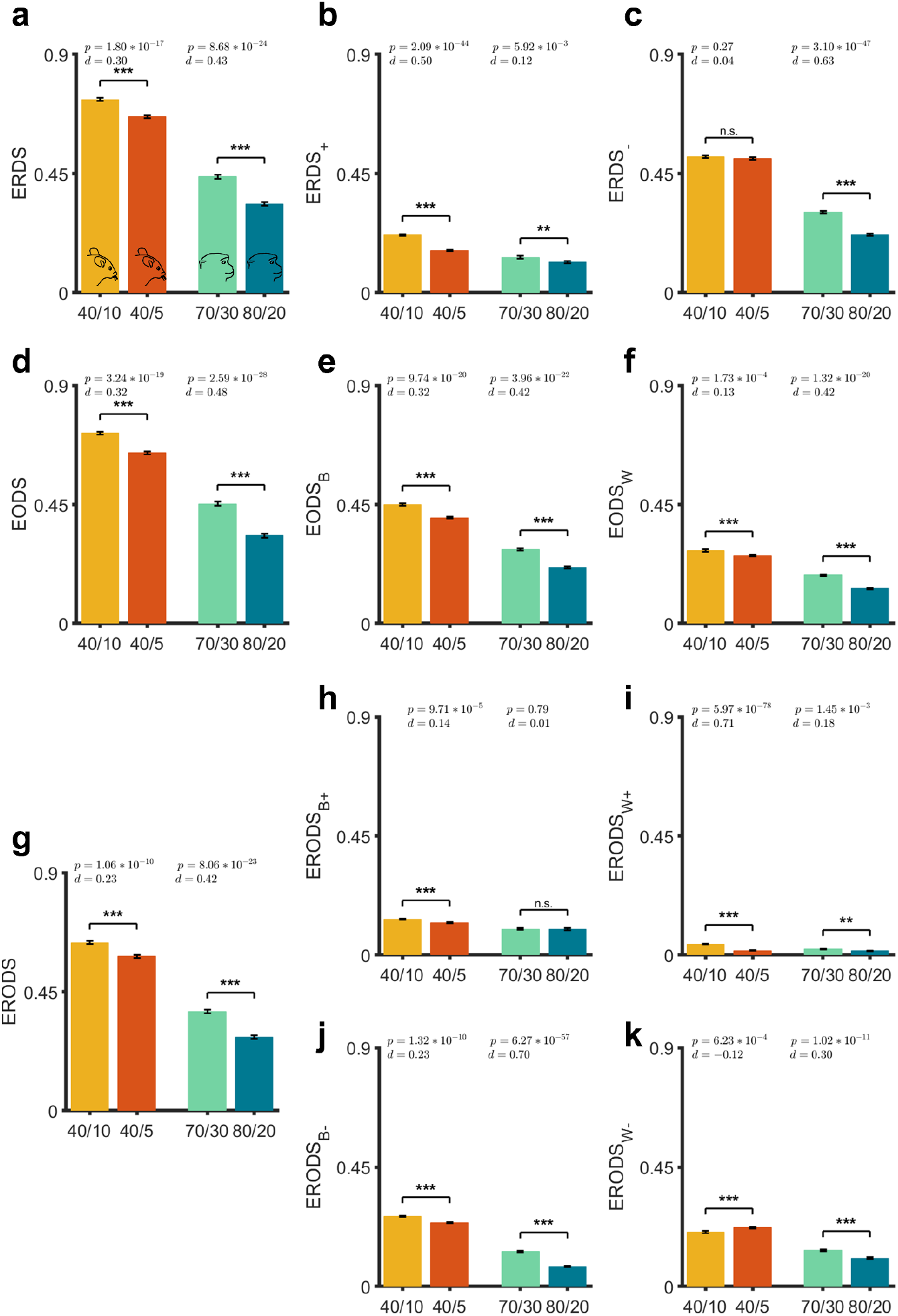
Entropy-based metrics capture changes in learning strategy between the two reward schedules. Plotted are ERDS and ERDS decompositions (a-c), EODS and EODS decompositions (d-f), and ERODS and ERODS decompositions (g-i) in the 40/10 and 40/5 reward schedules for mice and the 70/30 and 80/20 reward schedules for monkeys. Error bars indicate s.e.m. Reported are the *p*-values (two-sided t-test) and Cohen’s d-values. ERDS, EODS, and ERODS were significantly higher in the 40/10 schedule in mice and the 70/30 schedule in monkeys. Overall, increased entropy in the 40/10 schedule in mice and the 70/30 schedule in monkeys suggests a decrease in the consistency of reward and option-dependent strategy. Decreased consistency of reward and option-dependent strategy may be due to the greater similarity of reward probabilities for the two options in the 40/10 and 70/30 reward schedules.

**Figure S3.**
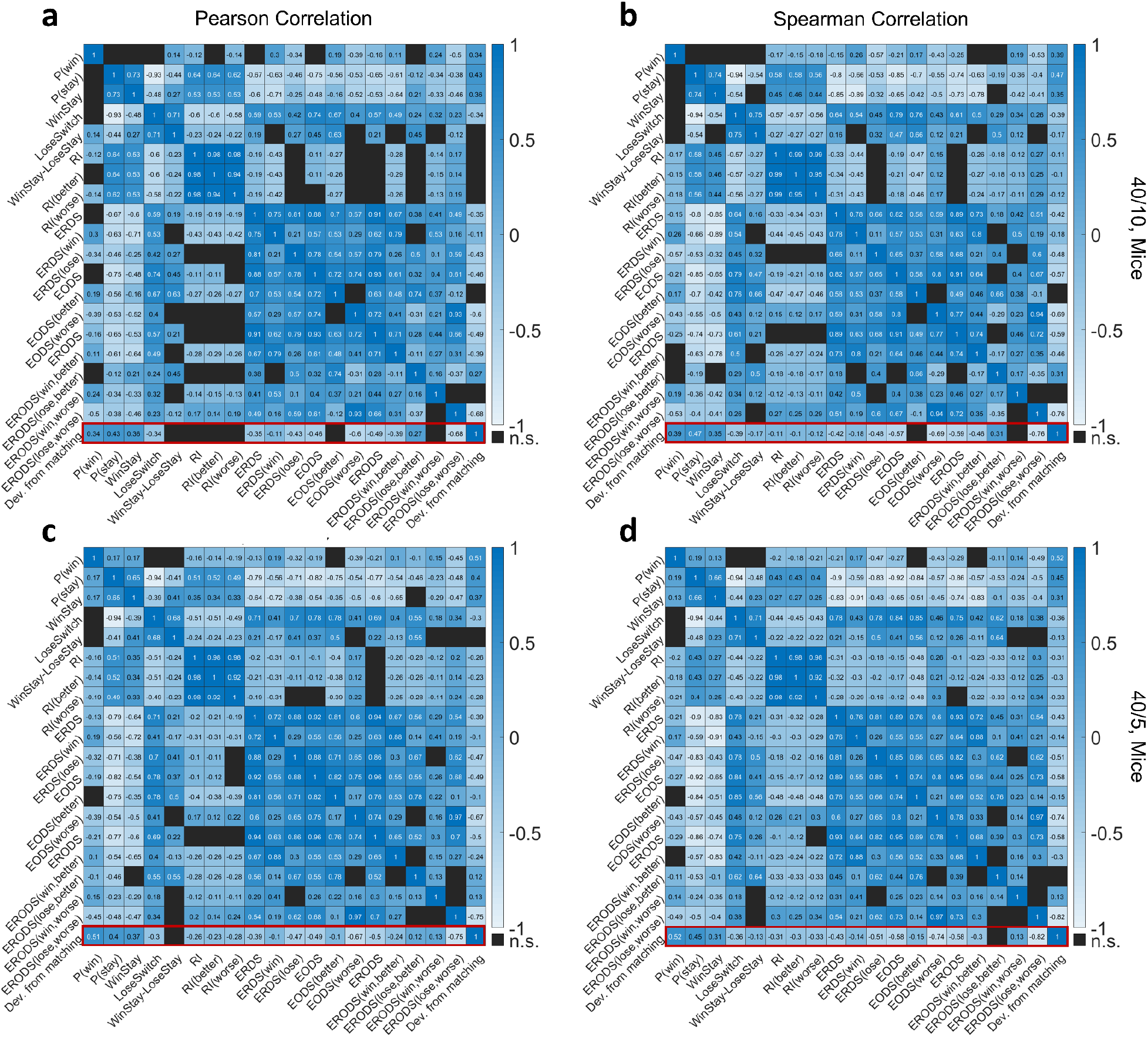
Correlation between undermatching and proposed entropy-based metrics separately for each reward schedule in mice. Correlation matrix for 19 behavioral metrics and undermatching using Pearson (a) and Spearman (b) tests, separately for blocks with reward probabilities equal to 40/10 (a-b) or 40/5 (c-d). Correlation coefficients are computed across all blocks, and matrix elements with non-significant values (*p* > .0001) are not shown (cells in black). The red rectangles highlight correlation coefficients between behavioral metrics and undermatching. Overall, the new entropy-based metrics were similarly strongly correlated with undermatching in both reward environments.

**Figure S4.**
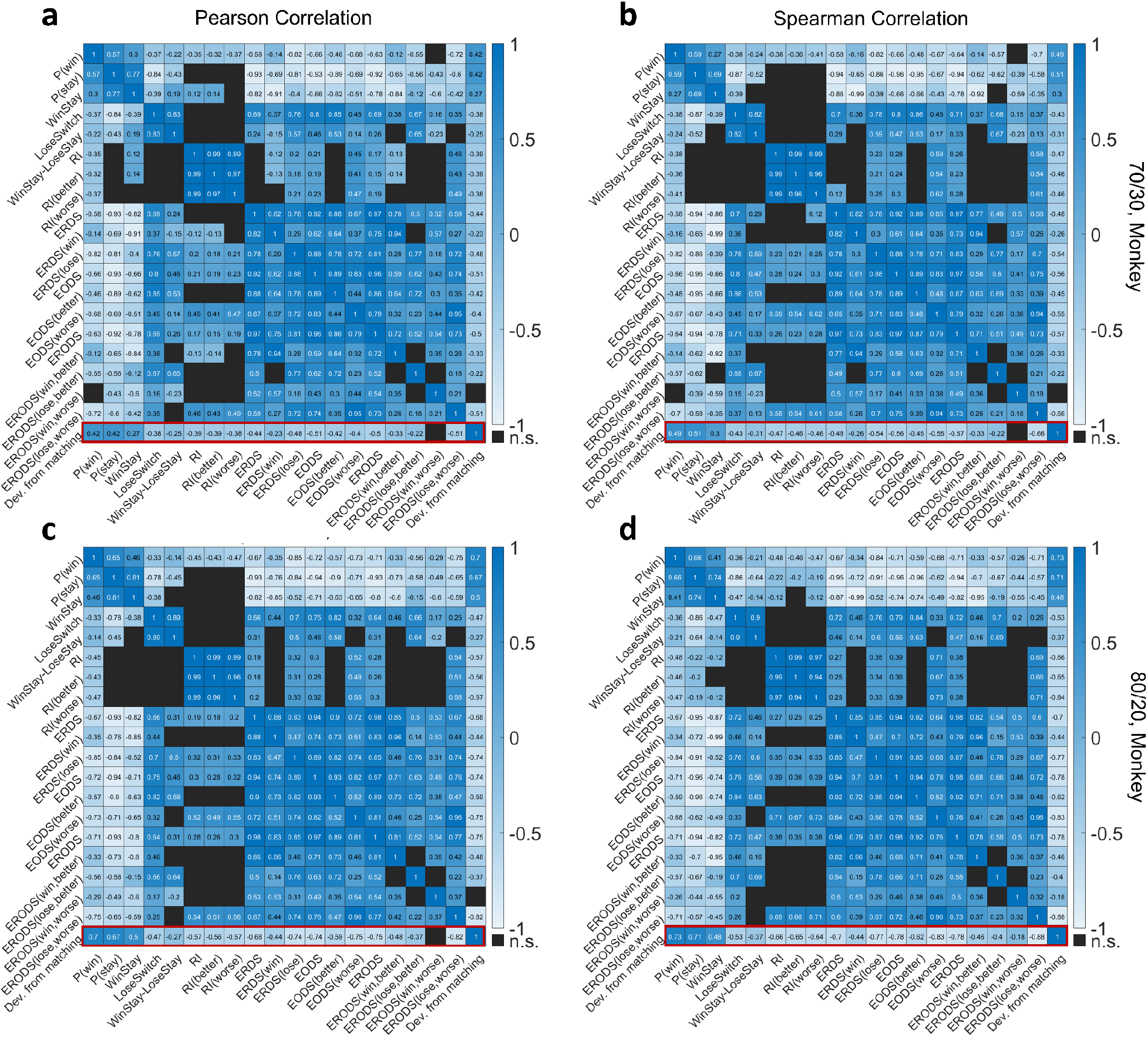
Correlation between undermatching and proposed entropy-based metrics separately for each reward schedule in monkeys. Same as Figure S3 but for monkeys, with blocks with reward probabilities equal to 70/30 (a-b) or 80/10 (c-d). Overall, the new entropy-based metrics were similarly strongly correlated with undermatching in both reward environments.

**Figure S5.**
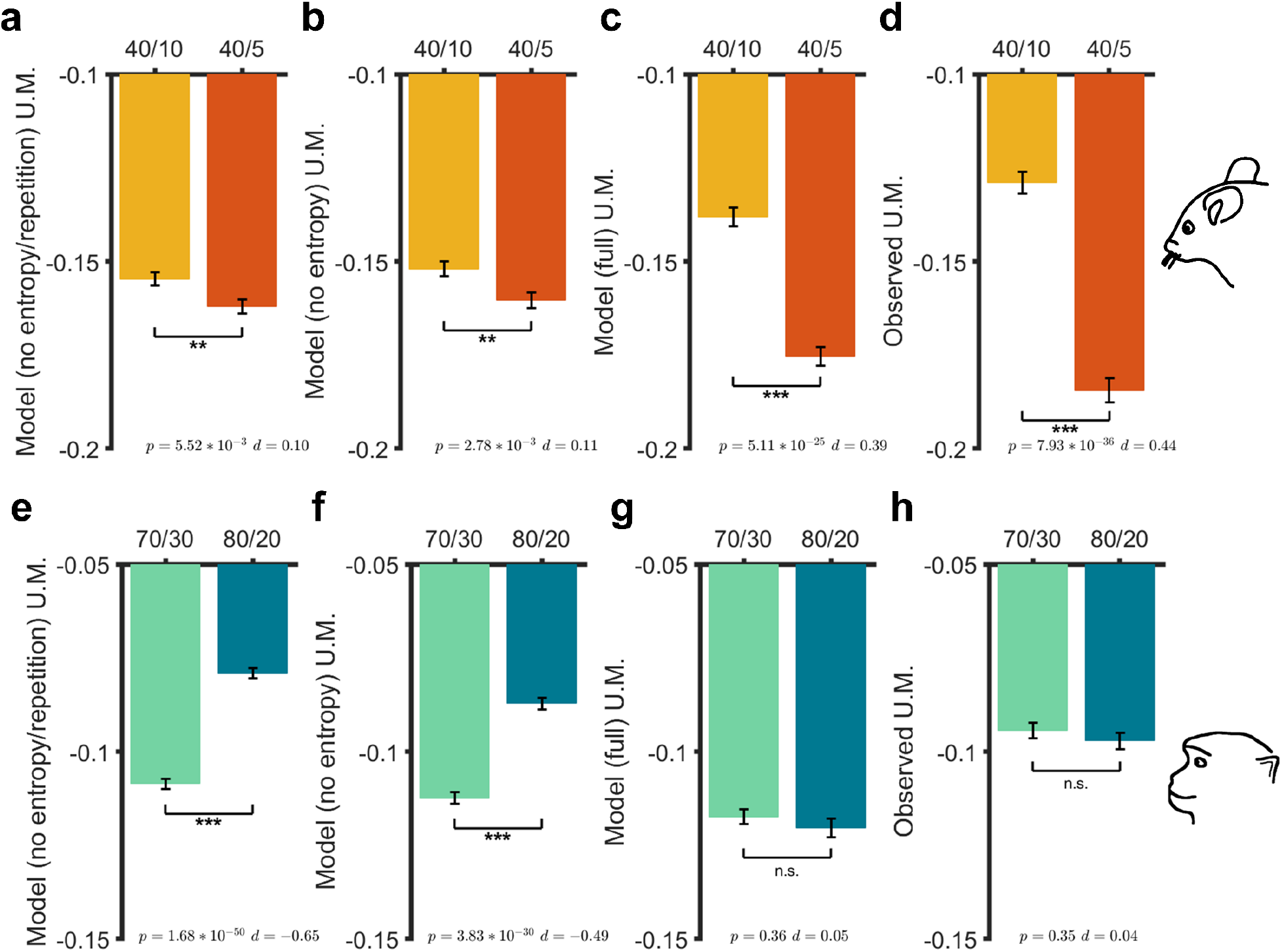
Entropy-based metrics capture differences in undermatching between reward environments. **(a–d)** Plotted are deviation from matching values in the 40/10 and 40/5 reward schedules for mice predicted from 10-fold cross-validated linear regression models using all behavioral metrics except entropy and repetition metrics as predictors (a), all behavioral metrics except entropy-based metrics as predictors (b), and all metrics as predictors (c) versus observed deviation from matching (d). Predictors were selected for inclusion using the stepwise regressions described in the manuscript, then 10-fold cross-validated linear regression was performed to fit models and predict undermatching (U.M.) (see **Methods**). Error bars indicate s.e.m. Reported are the *p*-values (two-sided t-test) on the left and Cohen’s d-values on the right. **(e–h)** Similar to (a-d), but with monkey data in the 70/30 and 80/20 reward schedules. The full regression model replicates the observed differences between reward schedules in mice and the lack of observed differences between reward schedules in monkeys.

**Figure S6.**
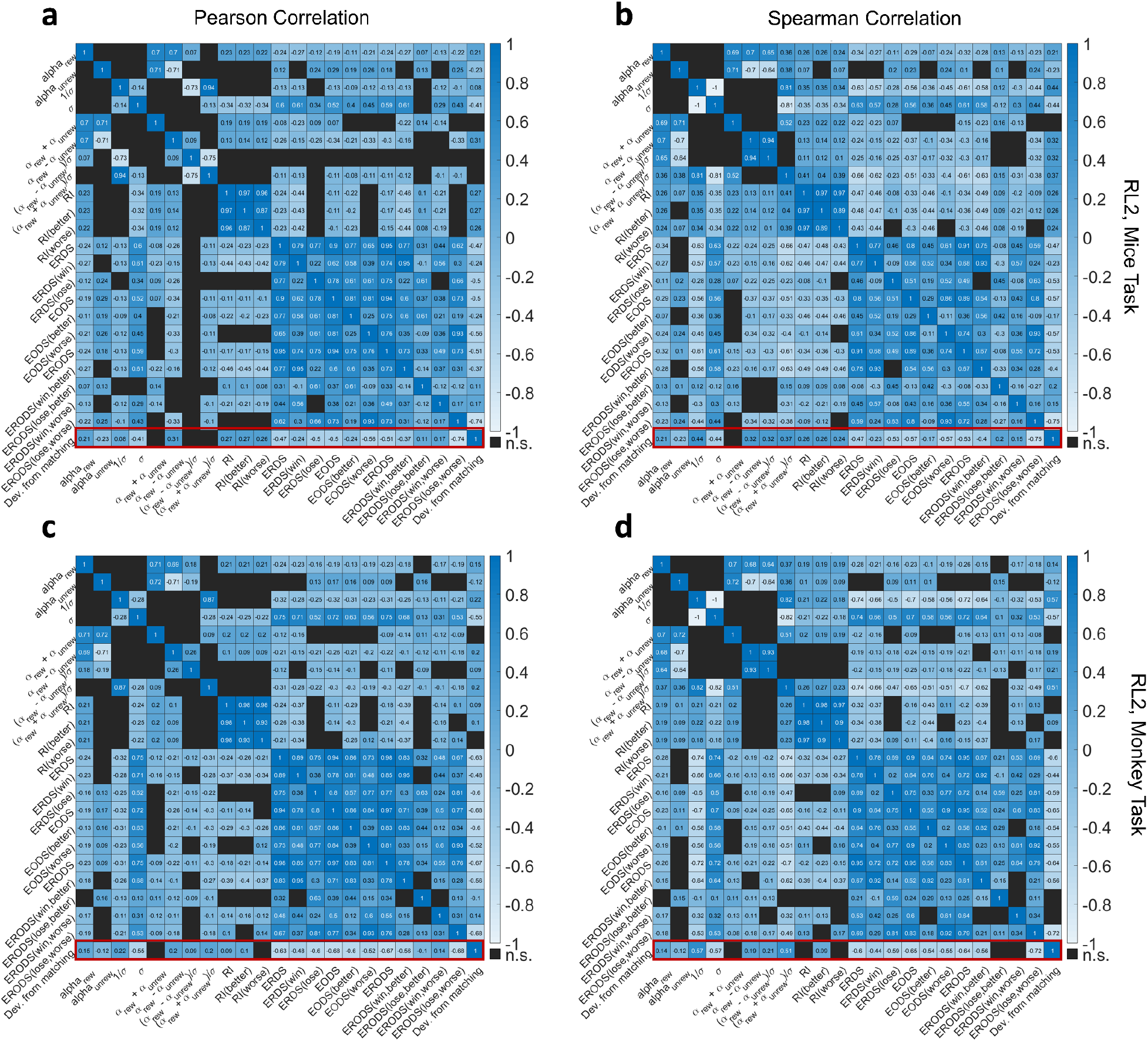
Undermatching in the RL model was better predicted by entropy-based metrics than parameters of the RL model. Plotted are correlation matrices for entropy-based metrics and deviation from matching computed over generated choice from block-wise simulations of RL2 with random parameters, separately based on Pearson (a,c) and Spearman (b,d) tests. Simulations were done using both the mice task setup and monkey task setup (a–b and c–d, respectively). Correlation coefficients are computed across all blocks, and matrix elements with non-significant values (*p* < .0001) are not shown (cells in black). The red rectangles highlight correlation coefficients between metrics and undermatching. Stochasticity in choice, σ, was the best correlate of deviation from matching out of all the RL parameters for both animals (*Pearson: mice: r* = −0.41, *monkeys: r* = −0.55; Spearman: mice: r = −0.44, *monkeys: r* = −0.57). In contrast, ERODS_W−_ was the best correlate of deviation from matching out of all entropy-based metrics (*Pearson: mice: r* = −0.74, *monkeys: r* = −0.68; Spearman: mice: r = −0.75, *monkeys: r* = −0.72). Interestingly, entropy-based metrics were also highly correlated with σ, suggesting that they capture explore/exploit decisions (see columns 3-4 of matrices). Overall, the entropy-based metrics predict deviation from matching better than the parameters of the RL models.

**Figure S7.**
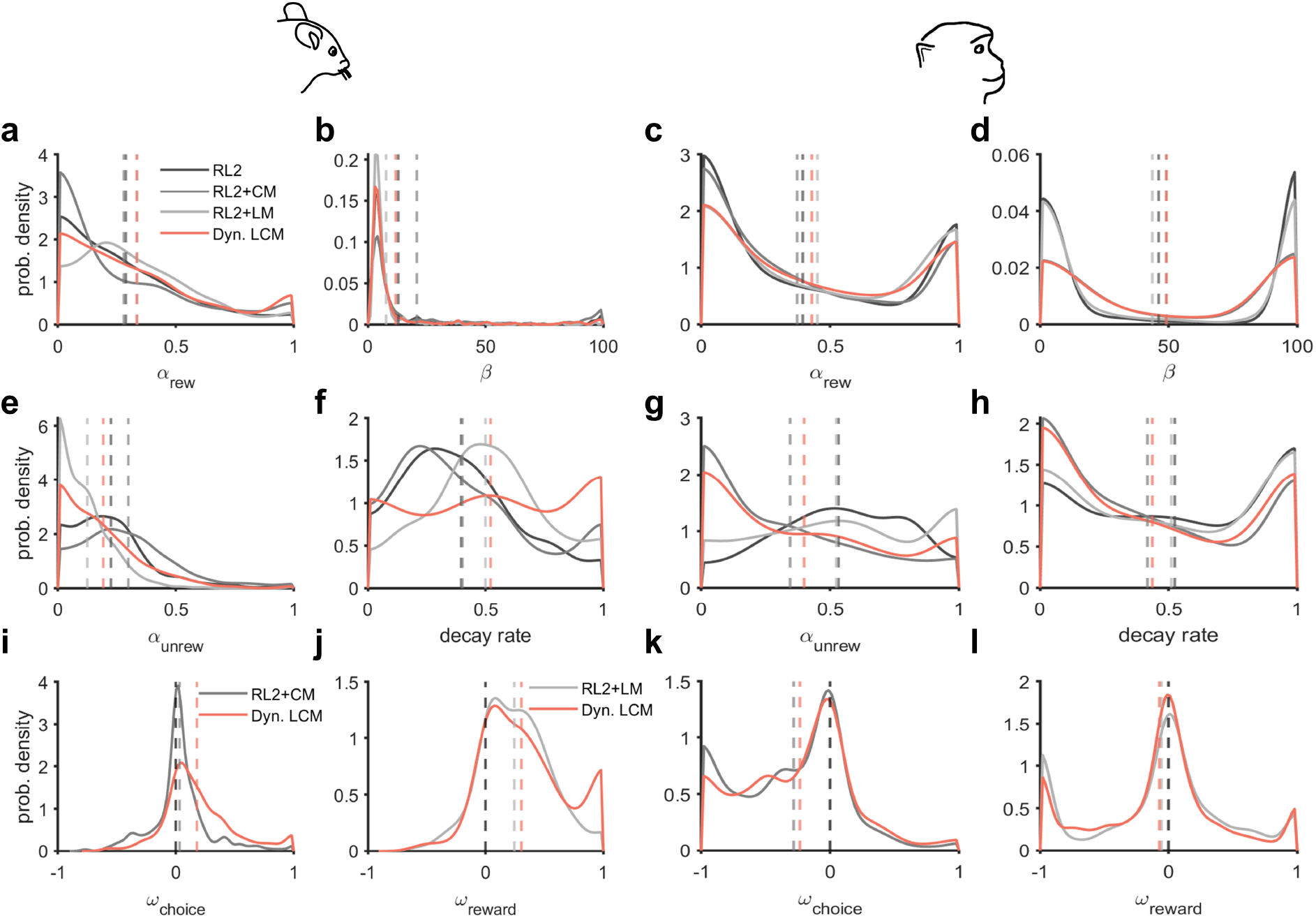
Distributions of model parameters obtained from fitting choice behavior. Plotted are distributions of the learning rate on the rewarded option (a,c), stochasticity of choice (b,d), learning rate on the unrewarded action/option (e,g), decay rate (f,h), the weight of the choice memory component (i,j), and the weight of the lose-memory component (k,l) for mice (a-j) and monkey (c-l) using four reinforcement learning models as noted in the legend. Distribution curves are estimated using kernel smoothing with reflection for boundary correction. Dashed vertical lines indicate the mean of each distribution in all plots and the black dashed vertical lines in (i-l) are zero lines. In mice, the choice and loss memory components have positive weights, whereas in monkeys the choice and loss memory components have slightly negative weights.

**Table S1.**
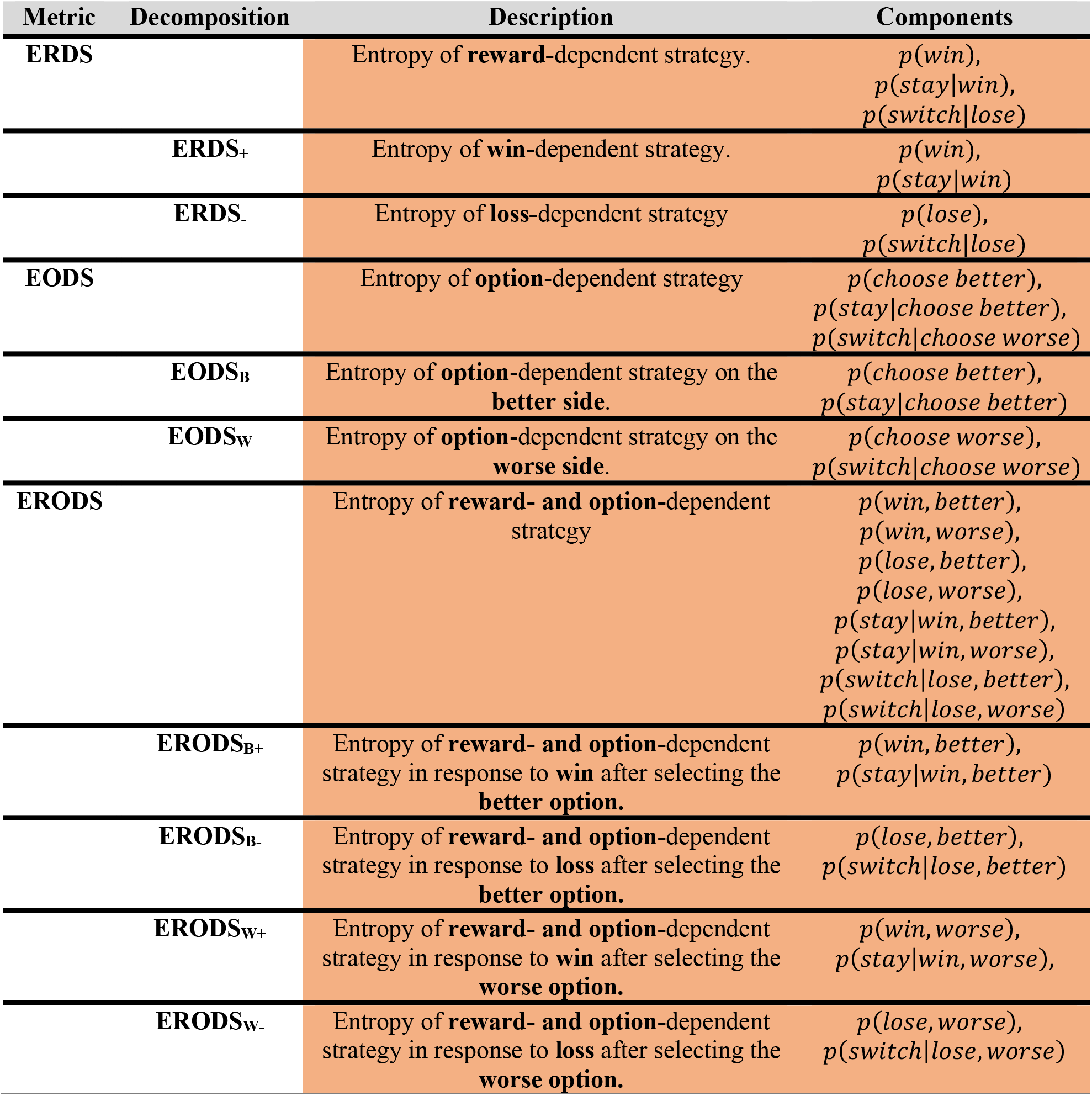
Summary and components of entropy-based metrics. Each row contains a short description of a metric, its decompositions, and the set of component probabilities that can be used to compute the metric.

**Table S2.** Various models used to fit choice data, their parameters and goodness-of-fit measures, and models’ ability to capture behavioral metrics. Each row provides a short description of a given model, its parameters, goodness-of-fit based on the negative log-likelihood values separately for mice and monkeys (column 4), McFadden *R^2^* values or variance in choice explained by each model (column 5), goodness-of-fit based on the AIC (column 6), D-values based on Kolmogorov-Smirnov tests comparing distributions of ERODSw- (column 7) and deviation from matching (column 8) predicted using model simulations and actual behavior. Values reported in parentheses and the asterisks in column 6 indicate the difference in AIC of a given model and the full model (Dyn. LCM) and the significance of this difference using paired-sample t-test (*p* < 3.0 × 10^-5^). Rows in orange and cyan correspond to mouse and monkey data, respectively.

| Model | Model Description | Parameters | -LL | MF $R^2$ | AIC | $D$<br>ERODSw- | $D$<br>matching |
| --- | --- | --- | --- | --- | --- | --- | --- |
| RL1 | Return-based RL | $\alpha_{\text{rew}}, \alpha_{\text{unrew}}, \beta$ | 228.31 | 0.184 | 462.62<br>(88.97*) | 0.238 | 0.224 |
|  |  |  | 21.05 | 0.620 | 48.11<br>(4.47*) | 0.054 | 0.092 |
| RL2 | Income-based RL | $\alpha_{\text{rew}}, \alpha_{\text{unrew}}, \beta, \text{decay}_{\text{rate}}$ | 189.61 | 0.322 | 387.22<br>(12.57*) | 0.121 | 0.091 |
|  |  |  | 18.09 | 0.674 | 44.18<br>(0.54*) | 0.072 | 0.101 |
| RL3 | Return-based RL + full choice kernel | $\alpha_{\text{rew}}, \alpha_{\text{unrew}}, \beta, \omega_{\text{CM}}, \gamma_{\text{CM}}$ | 190.72 | 0.318 | 391.44<br>(16.79*) | 0.095 | 0.088 |
|  |  |  | 18.24 | 0.671 | 46.47<br>(2.83*) | 0.027 | 0.072 |
| RL2+ CM | Income-based RL + choice memory | $\alpha_{\text{rew}}, \alpha_{\text{unrew}}, \beta, \text{decay}_{\text{rate}}, \omega_{\text{CM}}$ | 187.83 | 0.328 | 385.65<br>(11.00*) | 0.119 | 0.090 |
|  |  |  | 16.54 | 0.702 | 43.10<br>(-0.54*) | 0.037 | 0.065 |
| RL2+ CM+ | Income-based RL + choice memory with positive CM weight | $\alpha_{\text{rew}}, \alpha_{\text{unrew}}, \beta, \text{decay}_{\text{rate}}, \omega_{\text{CM}}$ | 188.37 | 0.326 | 386.74<br>(12.09*) | 0.122 | 0.090 |
|  |  |  | 17.90 | 0.677 | 45.79<br>(2.15*) | 0.072 | 0.101 |
| RL2+ LM | Income-based RL + loss memory | $\alpha_{\text{rew}}, \alpha_{\text{unrew}}, \beta, \text{decay}_{\text{rate}}, \omega_{\text{LM}}$ | 183.13 | 0.345 | 376.26<br>(1.61*) | 0.060 | 0.079 |
|  |  |  | 17.06 | 0.692 | 44.13<br>(0.49*) | 0.060 | 0.089 |
| RL2+ LM+ | Income-based RL + loss memory with positive LM weight | $\alpha_{\text{rew}}, \alpha_{\text{unrew}}, \beta, \text{decay}_{\text{rate}}, \omega_{\text{LM}}$ | 183.16 | 0.345 | 376.31<br>(1.66*) | 0.059 | 0.078 |
|  |  |  | 17.07 | 0.692 | 44.14<br>(0.50*) | 0.060 | 0.089 |
| Dyn. LCM | Income-based RL + loss memory + choice memory | $\alpha_{\text{rew}}, \alpha_{\text{unrew}}, \beta, \text{decay}_{\text{rate}}, \omega_{\text{CM}}, \omega_{\text{LM}}$ | **181.32** | **0.352** | **374.65** | **0.056** | **0.072** |
|  |  |  | **15.82** | **0.715** | **43.64** | **0.040** | **0.067** |

